# Nano-structured Hydrotrope-Caged Cytochrome c with Boosted Stability in Harsh Environments: A Molecular Insight

**DOI:** 10.1101/2023.02.05.527166

**Authors:** Pranav Bharadwaj, Dheeraj Kumar Sarkar, Meena Bisht, Sachin M. Shet, Nataraj Sanna Kotrappanavar, Veeresh Lokesh, Gregory Franklin, Jan Brezovsky, Dibyendu Mondal

**Affiliations:** Centre for Nano and Material Sciences, Jain (Deemed-to-be University), Jain Global Campus, Kanakapura, Bangalore, Karnataka, India – 562112; Laboratory of Biomolecular Interactions and Transport, Department of Gene Expression, Institute of Molecular Biology and Biotechnology, Faculty of Biology, Adam Mickiewicz University, Uniwersytetu Poznanskiego 6, 61-614 Poznan, Poland; International Institute of Molecular and Cell Biology in Warsaw, Ks Trojdena 4, 02-109 Warsaw, Poland; Institute of Plant Genetics (IPG), Polish Academy of Sciences, Strzeszyńska 34, 60-479 Poznań, Poland

**Author notes:** These authors have contributed equally to this work (Pranav Bharadwaj, Dheeraj Kumar Sarkar).

**Keywords:** ATP, Biological hydrotropes, Ionic liquid, Cytochrome c, Biocatalysis, Protein dynamics

## Abstract

Green and nano-structured catalytic media are vital for bio-catalysis to attenuate the denaturation tendency of biocata-lysts under severe reaction conditions. Hydrotropes with multi-faceted physiochemical properties represent promising systems for sustainable protein packaging. Herein, the suitability of adenosine-5’-triphosphate (ATP) and cholinium sa-licylate ([Cho][Sal]) ionic liquid (IL) to form nano-structures and to nano-confine Cytochrome c (Cyt c) were demonstrat-ed to enhance the stability and activity under multiple stressors. Experimental and computational analyses were under-taken to explain the nano-structured phenomenon of ATP and IL, structural organizations of nano-confined Cyt c, and site-specific interactions that stabilize the protein structure. Both ATP and IL form nano-structures in aqueous media and could cage Cyt c via multiple nonspecific soft interactions. Remarkably, the engineered molecular nano-cages of ATP (5-10 mM), IL (300 mg/mL), and ATP+IL surrounding Cyt c resulted in 9-to-72-fold higher peroxidase activity than native Cyt c with exceptionally high thermal tolerance (110oC). The polar interactions with the cardiolipin binding site of Cyt c, mediated by hydrotropes, were well correlated with the increased peroxidase activity. Furthermore, higher activity trends were observed in the presence of urea, GuHCl, and trypsin without any protein degradation. Specific binding of hy-drotropes in highly mobile regions of Cyt c (Ω 40-54 residues) and enhanced H-bonding with Lys and Arg offered excel-lent stability under extreme conditions. Additionally, ATP effectively counteracted reactive oxygen species (ROS)-induced denaturation of Cyt c, which was enhanced by the [Sal] counterpart of IL. Overall, this study explored the robustness of nano-structured hydrotropes to have a higher potential for protein packaging with improved stability and activity under extreme conditions. Thus, the present work highlights a novel strategy for real-time industrial bio-catalysis to protect mitochondrial cells from ROS-instigated apoptosis.

**Summary:** Suitability of ATP and [Cho][Sal] ionic liquid to form nanostructured hydrotropes and their utility in protein packaging in extreme conditions are discussed. Both ATP and IL form nanostructures in aqueous media and could cage Cyt c via multiple nonspecific soft interactions. The engineered molecular nanocages surrounding Cyt c resulted in 9-to-72-fold higher peroxidase activity than native Cyt c with exceptionally high thermal tolerance (110°C) and stability in the presence of urea, GuHCl, and trypsin without any protein degradation.

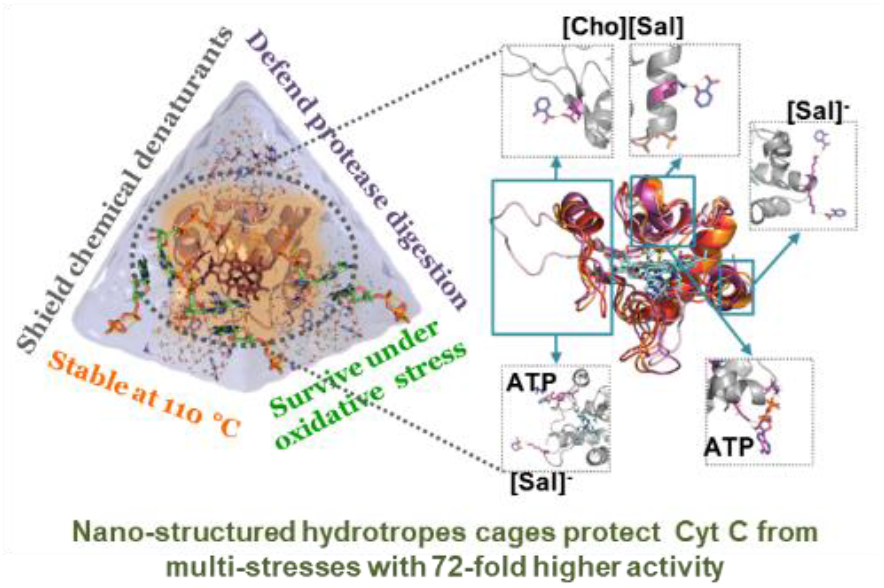

## Introduction

Adenosine-5’-triphosphate (ATP) has shown multi-faceted functions, including its participation in the electron transport chain, acting as the energy currency of cells, holding the key for caspase-9-activated apoptosis, and taking part in transcription by revamping chroma-tin complexes.^[1-3]^ The role of physiological concentrations of ATP in preventing thermally-induced protein aggregation in mammalian cells was previously demonstrated by Nguyen and Benusaude,^[4]^ and the answer regarding the regulation of a high concentration of ATP (2-10 mM) in the cell was unveiled by Patel and co-workers, whereby ATP is claimed as a biological hydrotrope.^[5]^ The term ‘hydrotrope’ was coined by Neuberg for amphiphilic molecules having a characteristic range of minimum hydrotrope concentration, which augment the solubility of partially soluble organic or hydrophobic substances in water.^[6-7]^ This opened a new channel to explore ATP in the medicinal field to treat Parkinson’s and Alzheimer-like diseases by preventing phase separation of biological fluids, which causes amyloid fibrillation.^[8-9]^ The solubilizing tendency of ATP was attributed to its nonspecific aggregation, charge reinforcement, and self-assembly through hydrophobic interactions (Figure 1a).^[10-11]^ Thus, the hydrotropic mechanism of ATP follows both the Neuberg and Hofmeister effects.^[12]^ Self-aggregated ATP is found to provide thermal stability to lysozyme, malate dehydrogenase, and ubiquitin,^[13]^ and so it is important to understand how the self-aggregation property of ATP is useful in protein packaging under multiple stressors, which has not been previously studied.

**Figure 1:**
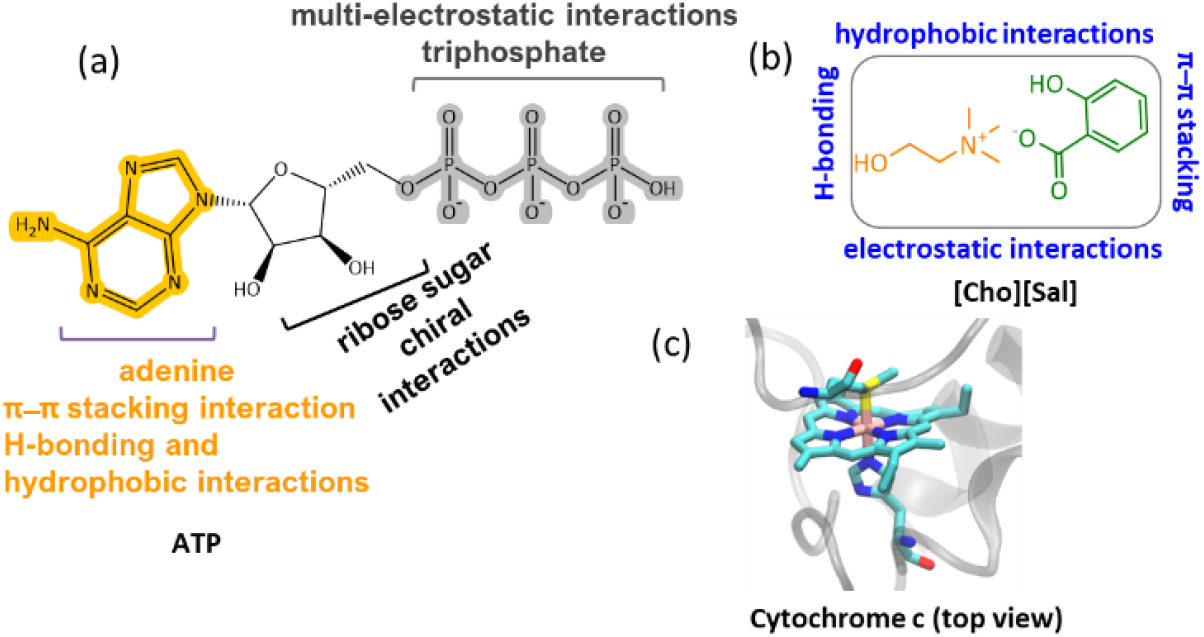
Chemical structures and structural features of (a) ATP and (b) [Cho][Sal] IL, with possible interaction sites. (c) The modeled active site structure of Cyt c, based on the crystallographic structure of bovine heart Cyt c (PDB code: 2B4Z) at 1.5 Å resolution.

Unlike ATP, various amphiphilic nano-structured solvents have recently emerged with promising applications in energy-efficient bioprocesses.^[14]^ Due to their highly manipulative nature, ionic liquids (ILs) have gained interest for their ability to increase enzyme solubility and thermal stability.^[15]^ Interestingly, bio-based ILs of the cholinium family have shown hydrotropic properties^[16]^ with greater hydrophobicity and can enhance the nano-structuring of amphiphilic solutes in aqueous media.^[17]^ Earlier reports have shown cholinium-based ILs for protein packaging with long-term stability and activity,^[15]^ but there is a lack of understanding of the underlying mechanisms by which hydrotropic ILs (potential amphiphilic molecules) can regulate protein stability and activity.^[18]^ In this regard, an IL with a salicylate counterpart caught our attention as it contains carboxylic and hydroxyl groups, along with a hydrophobic aromatic ring featuring the salicylate ion and showing multiple interaction sites in the aqueous medium (Figure 1b).^[19-20]^ Additionally, salicylic acid is an important phytohormone that alters the mitochondrial processes by amplifying the concentration of reactive oxygen species (ROS) and induces cells to undergo programmed cell death.^[21-22]^ During apoptosis, ATP plays a crucial role in caspase-9 activation, a process stimulated by cytochrome c (Cyt c) release from mitochondria.^[23]^ Since both ATP and salicylate are coupled to Cyt c, the latter was chosen as a model protein for this study (Figure 1c), and ATP and choline salicylate ([Cho][Sal]) IL as the nano-structured hydrotropes.

Cyt c is a mitochondrial-based 13 kDa monomeric protein comprising approximately 104 residues and is widely distributed across eukaryotes, bacteria, and archaea.^[24]^ Despite its significant role in electron transfer, this protein is of great interest in many technological applications like biosensors, synthetic receptors, and molecular imprinting among others.^[24]^ Cyt c also shows peroxidase-like activity and serves as a bio-catalyst in several catalytic transformations.^[24]^ The protein is highly vulnerable to denaturation under prevailing stress conditions such as extreme temperatures, oxidative stress, and proteolytic and chemical denaturants.^[25]^ Recently, a surface modification strategy using quantum dots, metal-organic frame-works,^[26]^ and DNA was reported to enhance the stability of Cyt c.^[25,27]^ However, the stability of the protein when confined within a hydrotrope system has not yet been reported. Considering the hydrotropism and probability of nano-structuring,^[24]^ the present study aimed to cage Cyt c in ATP and [Cho][Sal] IL-based hydrotropes, and thereby improve the activity and stability of the protein in gentle and harsh environments (such as high temperature, chemical denaturants, protease digestion, and oxidative stress). Therefore, detailed experimental and in silico approaches were undertaken to study the structure-function relationship of nano-confined Cyt c in ATP, [Cho][Sal] and ATP + IL at a molecular level to develop a novel and sustainable solvent manipulation strategy for dynamic protein packaging and robust bio-catalysis.

## Results and Discussion

### ATP and [Cho][Sal] IL as nano-structured hydrotropes

To understand the extent to which the IL affects the self-aggregation of ATP, which is vital for its role as a biological hydrotrope, we performed molecular dynamics (MD) simulations of 5 mM ATP, 300 mg/mL [Cho][Sal], and their mixture. The pair-wise interactions of ATP and [Cho][Sal] molecules in water and the mixture were investigated using radial distribution functions (RDF). As expected, ATP exhibited a self-aggregating tendency at 4-to-5 Å and up to 15 Å (Figure 2a). The self-aggregating propensity of ATP molecules from a chemical perspective in the presence of an IL can occur in three possible ways, first the anion-π interactions between the aromatic ring and negatively charged oxygen moiety of phosphate groups, secondly through a non-bonding aromatic ring interaction (H-bonding, π–π stacking interactions, and cation–π interactions) and third, H-bond formation between the sugar and triphosphate group.^[11]^ It was evident from our analysis that two ATP molecules preferentially formed stack-like configurations in water (Figure S1a). In the presence of [Cho][Sal], the IL-mediated H-bonding (Figure S1b) markedly decreased the occurrence of the stack-like configurations of ATP (Figure 2b). Notably, we found that the IL at a molecular concentration of 300 mg/mL forms dense the H-bonding network mediated by cholinium and salicylate (Figure S1c). Unsurprisingly, we have not observed any perturbation of the [Cho][Sal] nano-structure upon the addition of ATP, given its low concentration. In addition to RDFs, the self-aggregating behavior of ATP and ATP+IL was further verified by a concentration-dependent ζ-potential study (Figure 2c and Figure S2). At 1 mM con-centration, which is below the critical hydrotropic con-centration (2 mM), a single broad distribution peak was observed with maxima of -8.12 mV, indicating the existence of a single non-aggregated structure (Figure 2c). At 2.5 mM, the distribution curve shifted to a lower negative value with peak maxima of - 5.96 mV. This lowering in the ζ-potential indicates the beginning of aggregated structures similar to the self-aggregation of nano-particles as a result of the decrease in the electrophoretic mobility.^[28]^ This is further evidenced by the shift of maxima from -5.96 to -2.35 mV for 5 mM ATP. Finally, at 10 mM con-centration, a broad single peak with maxima at -1.42 mV was obtained, which indicates the presence of a self-assembled, nanostructured ATP state. Conversely, 200mg/mL of [Cho][Sal] shows a very broad Gaussian peak with a maximum at -9.12 mV (Figure S3), which shifts to a lower value of -4.02 mV in the presence of 5 mM ATP, justifying the hypothesis that the IL enhanced the self-assembly of amphiphiles.^[19]^ The hydrodynamic radius (R_h_) of 5 mM ATP was found to be 0.83 nm (slightly higher than R_h_ of monomeric ATP, which is 0.57-0.65 nm),^[10]^ suggesting the existence of nano-structured/oligomeric forms and supporting our earlier evidence. In the presence of the IL, the R_h_ of ATP increased to 1.12 nm (Figure S4).

**Figure 2:**
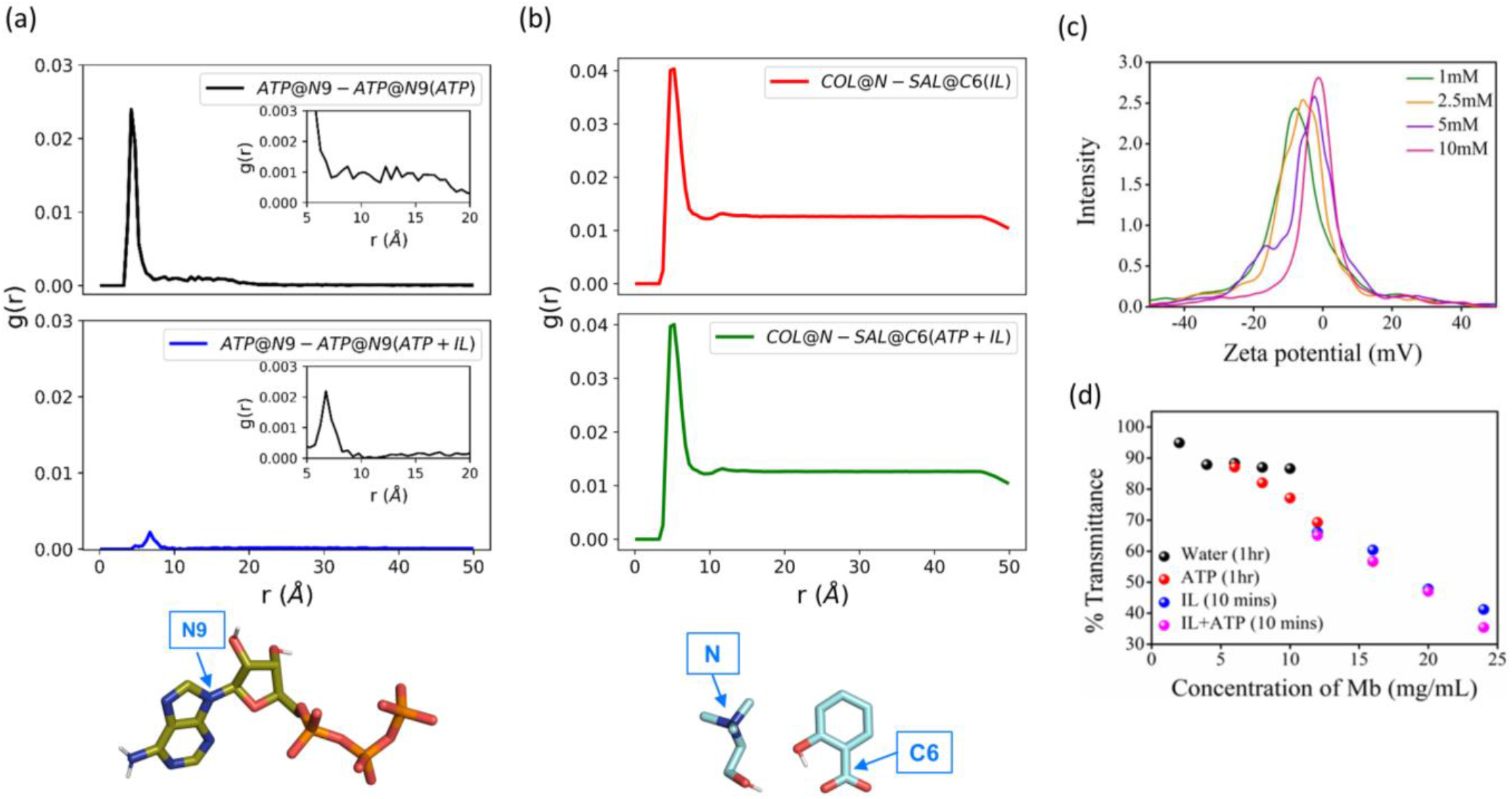
Mutual effects of ATP and [Cho][Sal] on their respective nano-structure and interactions. (a) RDF of 5 mM ATP (black) and 5 mM ATP in presence of [Cho][Sal] (blue). The N9 atom of ATP used for RDF calculation is shown by the blue arrow. (b) RDF of 300 mg/mL [Cho][Sal] (red) and [Cho][Sal] with added ATP (green). The N and C6 atoms used for RDF calculation are shown by the blue arrows. (c) Concentration-dependent Zeta potential analysis of ATP. (d) solubility data of equine heart myoglobin in an aqueous medium at pH 7 in the presence and absence of 5 mM ATP, 100 mg/mL [Cho][Sal], and a combination of both ATP+IL

To demonstrate the hydrotropic property of ATP and [Cho][Sal], solubility studies of myoglobin (Mb) were carried out in an aqueous medium at pH 7 in the presence and absence of 5 mM ATP, 100 mg/mL IL and a combination of both (Figure 2d). After 1hr, the maximum solubility of Mb was found to be 6 mg/mL. In contrast, ATP-induced solubility of Mb increased to 16 mg/mL during the same duration, which directly shows the hydrotropic nature of ATP. In the case of [Cho][Sal], the same solubility was achieved within 5 mins. Thus, the concentration of Mb was further increased and examined until 24 mg/mL. Interestingly, for ATP+IL, 24 mg/mL Mb showed higher solubility than the IL-only medium. These observations of enhanced solubility in the presence of two hydrotropes inspired us towards nano-confinement of Cyt c using ATP and [Cho][Sal]-based nano-structured hydrotropes to enable improved biological activity and stability of the protein.

### Structural features of nano-structured hydrotrope-caged Cyt c

To understand the structural features of caged Cyt c, four molecular systems were prepared, namely keeping Cyt c in water, ATP, IL, and both ATP + IL at 26.85°C (Figure 3a-d). The RDFs of hydrotropes in all systems with and without the presence of the protein were in agreement, indicating that the presence of Cyt c had a negligible effect on the adopted hydrotrope nano-structures (Figure 2a,b & Figure S5). A lower dielectric constant of the protein or more negative zeta potential of ATP and IL can be related to the colloidal stability of caged Cyt C.

**Figure 3:**
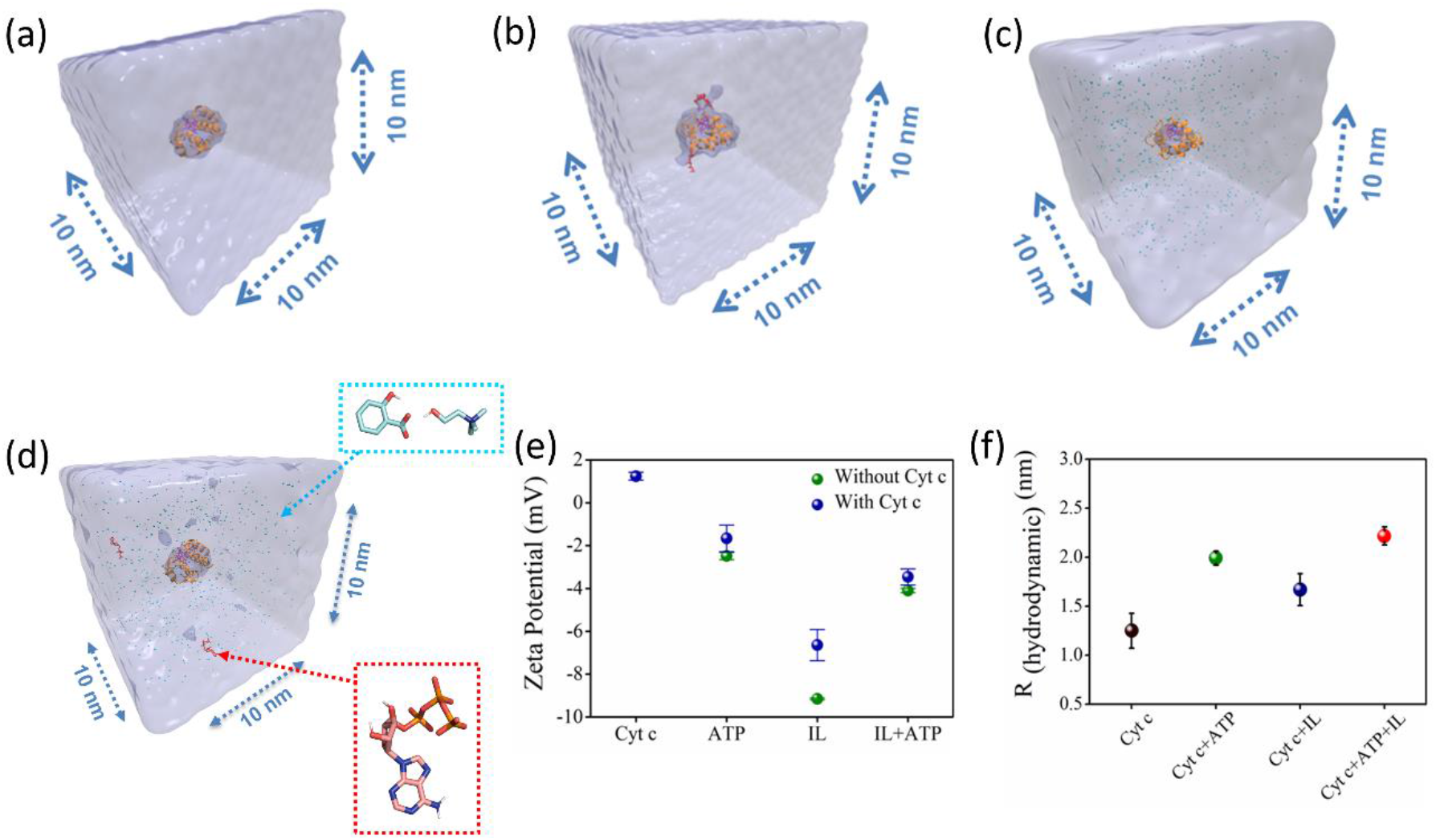
(a-d) Representative figures of a solvent of cubic box length 10 nm for Cyt c in H2O (a), ATP+ H_2_O (b), [Cho][Sal] + H_2_O (c), and ATP + IL + H_2_O molecular systems (d). Graph (e) shows zeta (ζ)-potential studies of nano-structured hydrotropes and Cyt c with and without nano-structured hydrotropes. Plot (f) depicts the hydrodynamic radius (R_h_) of native Cyt c and Cyt c caged in different nano-structured hydrotropes.

Accordingly, ζ-potential analysis was employed to envisage the electrostatic interaction between Cyt c and nanostructured hydrotropes. We found the ζ-potential of native Cyt c solution (10 μM) to be +1.24 mV (Figure 3e), and the ζ-potential of ATP (5 mM), IL (200 mg/mL), and ATP+IL was -2.48 mV, -9.16 mV, and -4.09 mV, respectively. After Cyt c incubation with ATP, IL, and IL+ATP, the ζ-potential values decreased to -1.66 mV, -6.64 mV, and -3.46 mV, respectively (Figure 3e). These changes in the ζ-potential demonstrate the possible electrostatic interaction between the protein and the nano-structured hydrotropes. At a lower concentration of Cyt c (2 μM), the ζ-potential of ATP increased from -2.48 mV to -4.38 mV, indicating the reorganization of a self-assembled ATP nanostructure around Cyt c (Figure S6).

Further R_h_ of native Cyt c was found to be 1.25 nm, which agrees with a previous report,^[30]^ and in the presence of ATP, the value increased to 1.99 nm showing self-assembly of ATP around the protein (Figure 3f and Figure S7). In the presence of IL+ATP, Rh of Cyt c is further increased to 2.22 nm. Thus, the hydrotropic mechanism of ATP occurs initially by charge reinforcement, followed by the stacking phenomenon. Along with ζ-potential and DLS, structures of nano-confined Cyt c were further characterized by UV-vis spectra (Figure S8). UV-vis spectrum of pure Cyt c comprises a Soret (B-band), Q band, and a shoulder peak (N-band), altogether representing the structural morphology around the heme group (Figure S8). In the presence of 5 mM ATP, a hyperchromic shift (a blue shift of 2 nm) in the Soret band (409 nm) was observed, suggesting that a non-polar environment dominates around the heme. In the case of IL (300 mg/mL [Cho][Sal]), absorbance in the Soret band enhanced considerably together with a blue shift, while the Q-band (520-550 nm) is broadened due to enhanced absorbance. This suggests changes in the tertiary conformations. An additional broad peak can be seen in the range of 620-695 nm, the intensity of which increases with an increased concentration of IL, demonstrating the local stability induced by [Cho][Sal].^[31]^ Moreover, the observed peaks suggest that a transit towards penta-coordinated Cyt c is formed in the presence of IL, leading to the formation of a high-spin Fe(III) heme center.^[31]^ A hypochromic shift in the Soret band region was observed for Cyt c caged within both ATP and IL nano-structures, indicating that the stability offered by both the hydrotropic systems by derived from driving the protein core towards the non-polar region. Overall, it is evident that the self-aggregation propensities of the hydrotropes persisted even in the presence of Cyt c, and the structure of caged Cyt c was partially affected (without any unfolding, as discussed below) due to multiple interactions with nano-structured hydrotropes.

### Nano-structured hydrotrope-caged Cyt c with boosted peroxidase activity: molecular insights on the improved catalytic activity and enhanced stability

In the presence of H_2_O_2_, Cyt c undergoes covalent modifications, which facilitate peroxidase-like activity.^[32]^ The activity of Cyt c in the presence of 5-10 mM ATP showed a similar trend with an >8-fold higher activity than native Cyt c (Figure 4a). Since a typical nano-structure is formed around Cyt c in the range of hydrotropic concentration (5-10 mM), similar relative activity is achieved rather than a maxima-minima trend. ATP acts as a hydrotrope initially by unwinding the protein and then stabilizing the extended chain through electrostatic interactions,^[11]^ which may be a reason for the high peroxidase activity. In the case of the [Cho][Sal] IL, nearly the same relative activities were seen from the 200-to-500 mg/mL range (Figure 4b). A clear trend was observed with the highest relative activity of 67-fold at the concentration of the IL of 300 mg/mL. The activity data agree with the UV-vis absorption spectra that inferred the transition of Cyt c towards penta-coordinated high spin complex. When the Met80 ligand is displaced, Cyt c naturally exhibits pronounced peroxidase-like activity.^[32]^ Interestingly, a replica of the IL optimization trend pat-tern was observed when using the ATP+IL mixture when [ATP] was kept constant (5 mM) and IL was varied (Figure 4c). At 300 mg/mL IL and 5 mM ATP concentrations, the activity of Cyt c was enhanced 72-fold more than native Cyt c. This activity enhancement is indicative that the structure of Cyt c is still intact and combined hydrotropic systems boost the peroxidase activity further. Some ILs enhance the self-assembly of co-solutes in the media due to the virtue of their solvophobic effects.^[17]^ Accordingly, the concentration of ATP was varied with IL being constant (300 mg/mL). A similar trend was observed with a >69-fold increase in activity (Figure 4d). These trends demonstrate that both IL and ATP further enhance the conformational changes of Cyt c, followed by stabilization of the extended protein structure by the self-assembling of ATP to surround it. Similar inference can be observed from Figure S5a (blue line), in which ATP tends to self-aggregate, even in the presence of IL and Cyt c.

**Figure 4:**
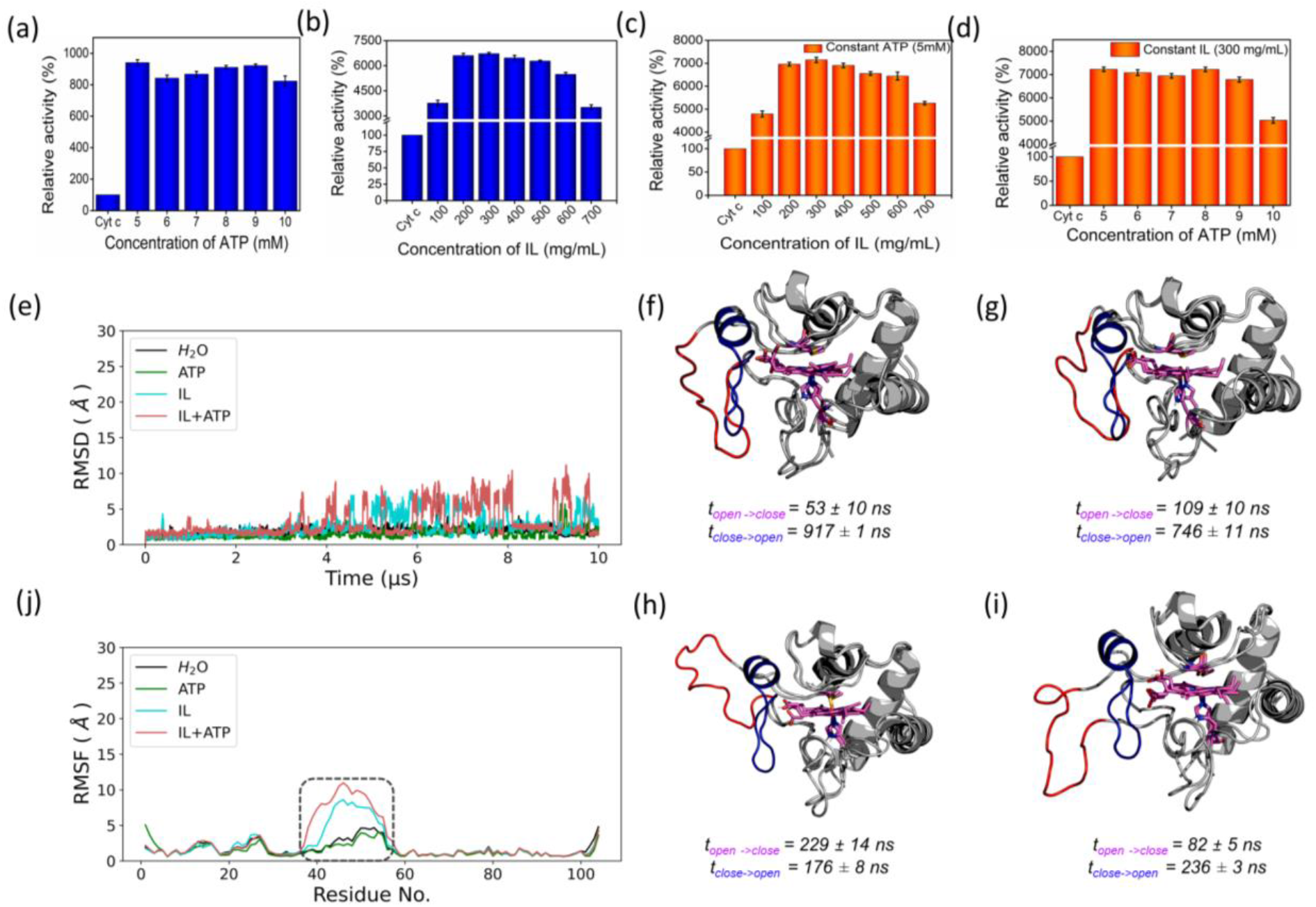
Peroxidase-like activity of Cyt linked to its dynamics. Optimization results of peroxidase-like activity of Cyt c with different hydrotropic concentrations of (a) ATP, (b) [Cho][Sal] IL, (c) varying [IL] by keeping [ATP] constant (5 mM), and (d) varying [ATP] by keeping IL constant (300 mg/mL). Structural characterization of Cyt c in four solvent systems at 26.85°C. (e) Stability and dynamics of Cyt c structure simulated in four solvent systems shown as backbone RMSD and RMSF, respectively. The peaks in the RMSD plot correspond to the reversible openings of Ω 40-54 residue region highlighted in the RMSF plot by a dashed rectangle. Figures (f-i) show the metastable states representing open (red) and closed (blue) structures of Cyt c in H_2_O, ATP, IL, and ATP+IL, respectively. The timescales of opening and closing processes were calculated as mean first passage times between the two states presented as mean ± standard deviation.

To understand the behavior of nano-structured hydrotrope-caged Cyt c, adaptive sampling MD simulations with a total of 10 μs simulation time for each system were performed. In all systems, we observed reversible structural rearrangements of the protein backbone (Figure 4e), which were more pronounced in mixtures containing IL and ATP+IL. Next, we traced these changes to the reversible opening of the loop in the Ω 40-54 residue region that was relatively more mobile than other parts of the protein for all four systems (Figure 4j). This region corresponds to the mAb 1D3 binding site and is reported to be more dynamic,^[33]^ while having specificity towards Cytochrome c oxidase.^[34]^ To delineate the details of this process and its modulation by the action of nano-structured hydrotropes, we have constructed Bayesian Markov state models from the simulations. This inference revealed the presence of metastable states with an open conformation of the Ω 40-54 loop of Cyt c in all systems (Figures 4f-i), which were characterized by their increased RMSD and elevated radius of gyration (Figure S9). The interactions of ATP and IL with Cyt c resulted in marked effects on the stability and dynamics of this region. In the presence of ATP, the closing process was approximately 2 times slower, whereas the opening became approximately 20% faster (Figure 4g). Similarly, the presence of the IL resulted in a much faster opening process and the closing speed was slowed even more prominently than that observed for ATP (Figure 4h). Finally, the dynamics of Cyt c in the mixture of ATP and IL exhibited ATP-like closing and IL-like opening times, enhancing the dynamics of this functionally important region (Figure 4i).

Since both ATP and IL were significantly affecting the dynamics of Cyt c, we identified the residues exhibiting the most frequent H-bonding with hydrotropes in the simulations (Figure 5). In general, ATP interacted preferentially with Lys residues (Figure S10a), which is in accordance with an earlier report demonstrating that triphosphate group of ATP can deliver weak, nonspecific interactions with conserved Lys or Arg residues^.[10]^ Additionally, the IL had a wider range of residues as relevant binding partners, although Lys was the most populated in terms of H-bonding (Figure S10b).This trend was also maintained in the mixture of ATP and IL (Figure S10c). We observed marked overlap in particular interaction hot spots for IL and ATP on the surface of Cyt c (Figure S10d,e). ATP most often formed interactions with K13, K25, K27, K72, K86, and K87 (Figure S11a,c). Notably, K72 is involved in site A of Cyt c, which is crucial for interacting with cardiolipin,^[35]^ and was reported to have a significant role in the peroxidase activity of Cyt c.^[36]^ Also, IL demonstrated general preference to Lys, with somewhat higher affinity towards K7, K79, R38 (Figure S11b,d). Interestingly we also found considerable affinity towards Y97, which has been ascribed a significant role in keeping the terminal helix intact and hence providing structural stability to the protein.^[37]^

**Figure 5:**
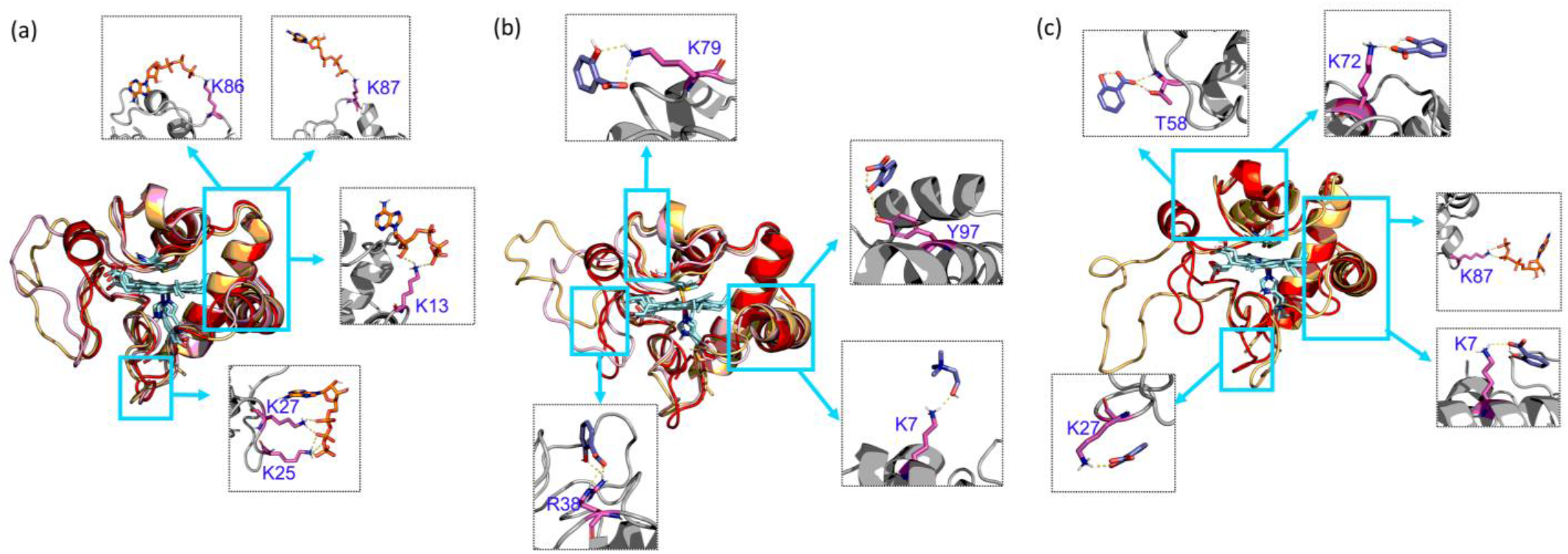
Interaction of hydrotrope molecules in metastable states in the open, closed, and transient Cyt c conformations from 10 μs of adaptive simulation data with (a) ATP, (b) [Cho][Sal] IL, and (c) ATP + IL. The top residues that have hydrogen bond interactions between solvent molecules and Cyt c are shown in gray squares (dashed), and the protein regions are highlighted in cyan (solid). All simulations were performed at 26.85°C. In all cases, the concentration of ATP and IL was 5 mM and 300 mg/mL, respectively.

### Nano-structured hydrotrope-caged Cyt c with enhanced thermal stability for high-temperature bio-catalysis

The activity of Cyt c increases with temperature up to 70°C, above which the protein starts to denature gradually (Figure 6a). Above 80°C, the activity drops due to significant denaturation of the protein.^[25]^ A similar trend is observed in the presence of ATP, but the activity extends to approximately a 20-fold increase than for native Cyt c at 70°C. In the case of IL and ATP+IL systems, the denaturation is prevented even up to extremes of 110°C. At 90°C, both Cyt c+IL and Cyt c+IL+ATP showed an accelerated activity value corresponding to 76-80-fold higher than native Cyt c. The stability aspects of Cyt c, when coupled to hydrotropes at elevated temperatures, were supported by UV-vis spectra (Figure 6b). Enhancement in the Soret band intensity was observed for both IL and IL+ATP at 90°C compared to the results obtained at 26.85°C, which implies that the secondary conformation, formed due to weak interactions between the hydrotrope systems and Cyt c, was preserved. ATP+IL showed an intensified Soret peak compared to the other conditions at 110°C, suggesting the advantage of engineering Cyt c with two different hydrotropes when aiming for high-temperature bio-catalysis. Additionally, as evident from non-reducing SDS-PAGE (Figure 6c), improved activity trends at high temperatures were achieved without structural degradation of the protein.

**Figure 6:**
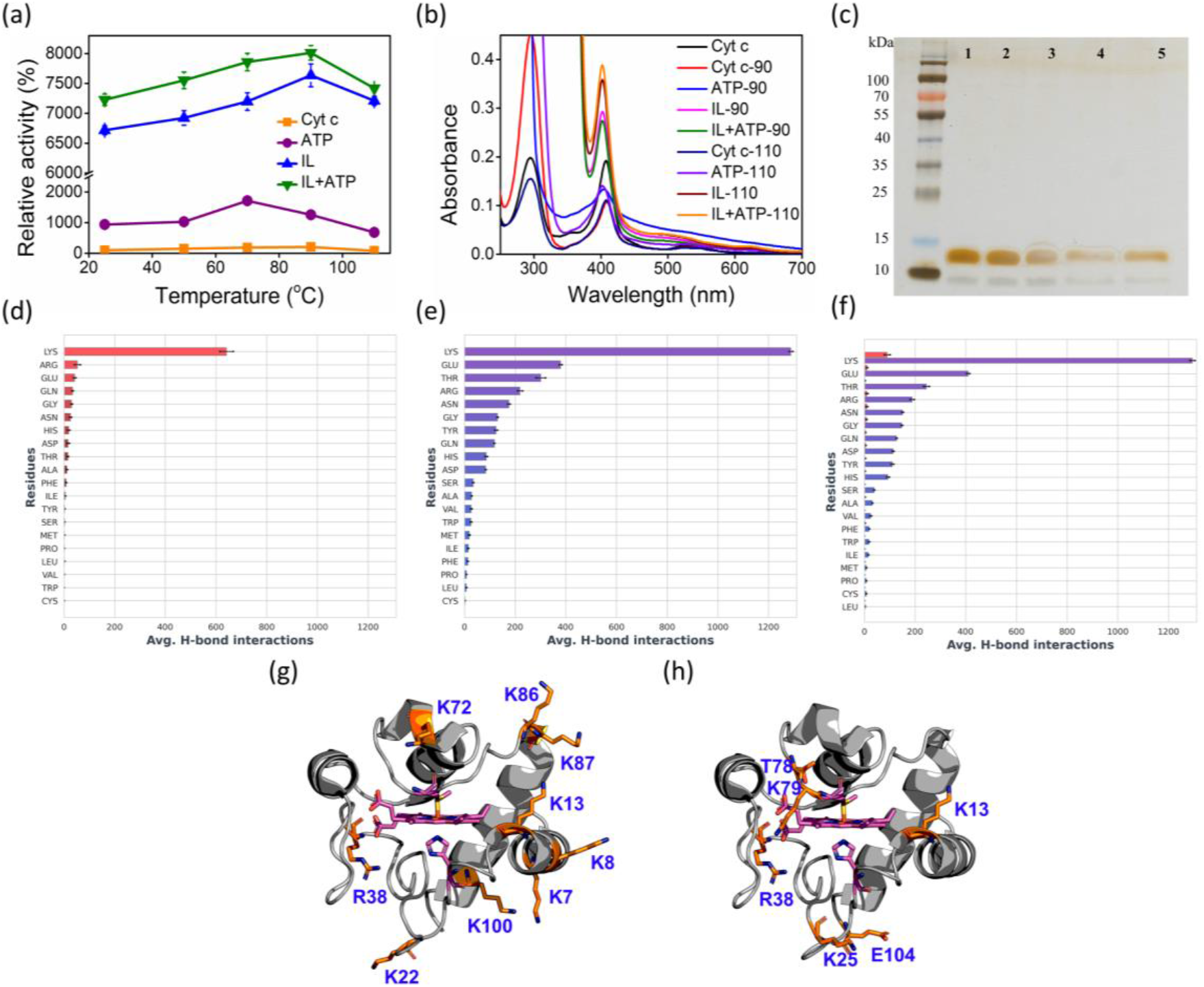
(a) Peroxidase activity of Cyt c with and without nano-structured hydrotropes at different temperatures. (b) UV-vis spectra of Cyt c at higher temperatures (90°C and 110°C) with and without nano-structured hydrotropes. (c) SDS-PAGE image of native Cyt c (lane 1), Cyt c incubated at 100°C (lane 2), Cyt c + ATP incubated at 100°C (lane 3), Cyt c + [Cho][Sal] incubated at 100°C (lane 4), and Cyt C + ATP + [Cho][Sal] incubated at 100°C (lane 5). (d-f) H-bonding interaction counts of individual residue types obtained from adaptive simulation data of Cyt c in the presence of ATP (d), IL (e), ATP and IL in the ATP+IL system (f) are depicted with ATP in red and IL in blue. All simulations were performed at 90°C. (g, h) The position of residues having a higher occurrence of H-bonding with ATP (g) and IL (h) also in their mixtures, respectively. The concentration of ATP and IL was 5 mM and 300 mg/mL, respectively.

A systematic computational study was undertaken at 90°C to gain structural insights into the higher stability of nano-structured-caged Cyt c at elevated temperatures. As expected, the Cyt c conformational ensemble was significantly altered at the higher temperature, with backbone RMSD being approximately 3-4 times higher than that at 26.85°C for all four solvents (Figure S12a). No significant effects were observed between the mobility of the Ω 40-54 residue region in the four systems than that observed for 26.85°C (Figure 4e and S12b). However, the thermal stability of the Ω 70-85 residues region was significantly enhanced with nano-structured hydrotropes compared to water (Figure S12b). From Bayesian Markov state models constructed based on the high-temperature simulations (Figure S12c-f), we could observe that Cyt c in water exhibited severe conformation-al instability, underlined by a very high RMSD of the open state, as well as the presence of multiple secondary open-like states (Figure S13). In contrast, the presence of ATP or/and IL approximately halved the maximal range of displacements of the Cyt c backbone, providing heightened Cyt c resistance. Hence, it is evident that ATP and IL could thermally stabilize the functional component of Cyt c via multiple polar interactions.^[24]^ Affinity of hydrotrope molecules towards specific amino acids at the higher temperature (Figures 6d-h and Figure S14) was found to be similar to that observed at 26.85°C (Figures S14 & S11). Overall, Lys and Arg were found to have frequent H-bonding at 90°C to ATP (Figures 6d, f,g). Similarly to the lower temperature environment, the IL had a broader range of specificity, and more affinity with Lys, Glu and Thr in terms of H-bonding (Figures 6e,f,h). The highest affinity was seen towards K7, K8, K13, K22, R38, K72, K86, K87,and K100 with ATP (Figure 6g), and K13, K25, R38, T78, K79, and E104 with IL (Figure 6h), all of which are crucial in maintaining thermal stability.^[38]^

### Improved stability of nano-structured hydrotrope-caged Cyt c when exposed to chemical denaturants

Above a particular concentration and on more pro-longed exposure to some chaotropic chemicals like H_2_O_2_, guanidine hydrochloride (GuHCl), and urea cause amide bond disruption or overexpose the metallic core of the enzyme, causing a loss of structural integrity and function. To shield the Cyt c against such chemical denaturants, the protein was incubated with the three nano-structured hydrotropic systems for 10 mins, followed by the addition of a chemical denaturant. After the interaction with 6 M GuHCl for 15 mins, there was a complete disruption in the heme pocket of native Cyt c and the activity severely declined, with the remaining activity approximately ∼20%. Conversely, hydrotrope-caged Cyt c maintained initial activity of 8.5-fold, 63-fold, and 66-fold in the presence of ATP, IL, and ATP+IL, respectively (Figure 7a). While urea is known to be a hydrotrope, it is well recognized as a chemical denaturant at concentrations above 6 M. In the pre-denaturation concentrations of urea (1-6 M), a ∼30 fold increase in peroxidase-like activity was obtained for Cyt c.39 After 30 mins incubation with 8 M urea, its activity was recorded at approximately 70% of the initial activity, while IL and ATP+IL hydrotrope-stabilized Cyt c showed similar activity (Figure 7b).

**Figure 7:**
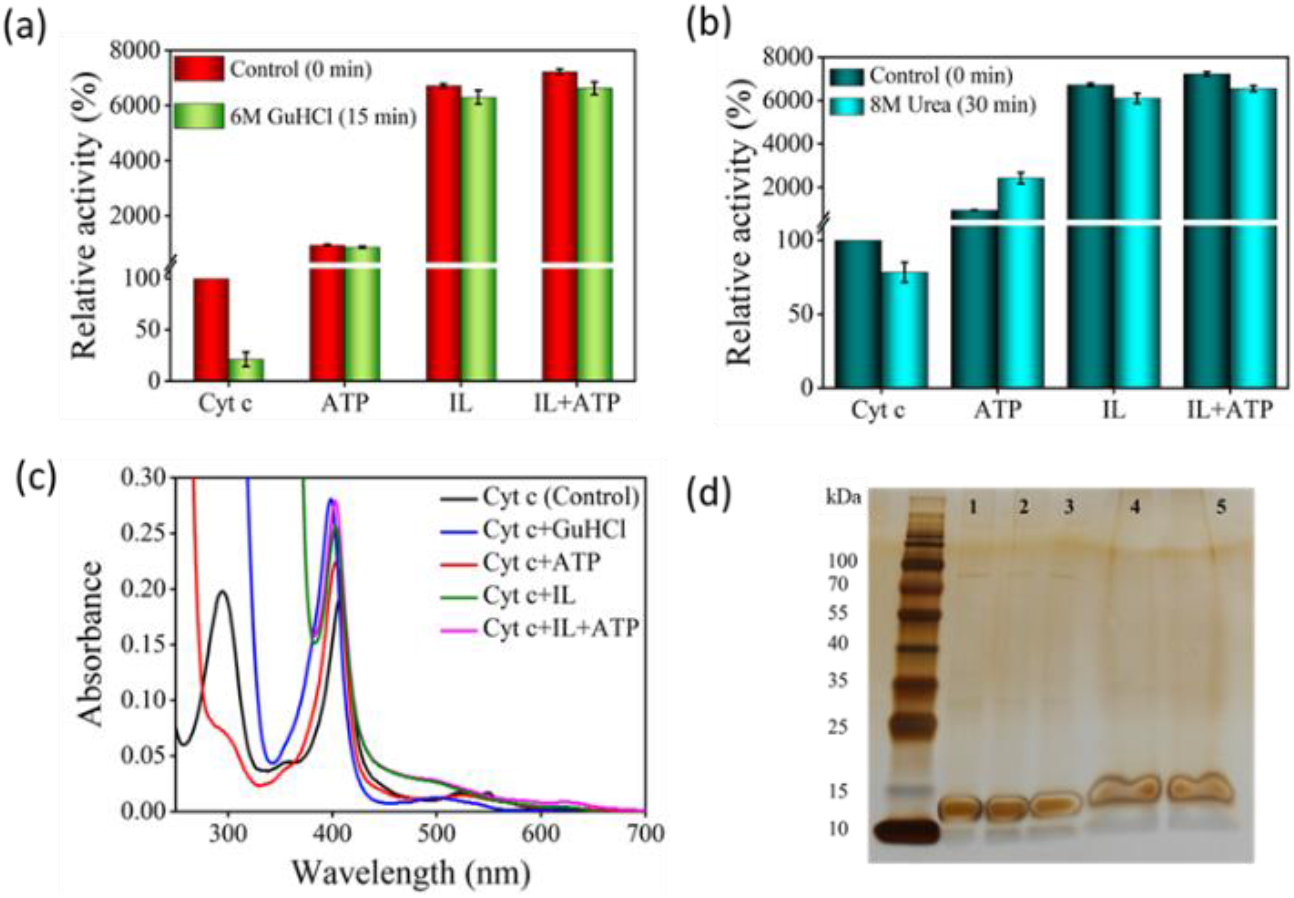
Effect of chemical denaturants, 6 M GuHCl (a) and 8 M urea (b) on the peroxidase activity of Cyt c and nano-structured hydrotrope-caged Cyt c. (c) UV-vis spectra of Cyt c after incubation with 6 M GuHCl. (d) SDS-PAGE image of native Cyt c (lane 1), Cyt c incubated with 8 M urea (lane 2), Cyt c + ATP incubated with 8 M urea (lane 3), Cyt c + [Cho][Sal] incubated with 8 M urea (lane 4), and Cyt c + ATP + [Cho][Sal] incubated with 8 M urea (lane 5). In all cases, the concentration of ATP and IL was 5 mM and 300 mg/mL, respectively.

Native Cyt c showed a denaturing pattern in the UV-vis spectra with GuHCl (Figure 7c), where the Q band structure was entirely lost and the Soret band was severely blue-shifted. In the case of ATP, changes in the 280 nm shoulder peak suggest partial dissociation of aromatic residues like Tyr and Trp. In contrast, the structures remained almost the same in the cases of IL and ATP+IL, except that the absorbance of the IL-based system trended lower. In the case of ATP, the activity of Cyt c almost doubled (∼24-fold higher than native Cyt c) in the presence of urea compared to the Cyt c+ATP system (Figure 7b). This strange observation suggested that ATP cannot prevent the further unfolding of Cyt c in the presence of urea, a behavior con-firmed by the UV-vis spectra. For native Cyt c in water, a blue shift similar to GuHCl along with the disruption of the Q band was observed, but for the Cyt c-ATP system, the Q band was retained. The hypochromic shift in the Soret peak for Cyt c-ATP indicated the beginning of structural deformation. IL and ATP+IL followed the regular trend, and this lower activity can be envisioned by the reduced Soret peak intensity (Figure S15). Further-more, SDS-PAGE analysis demonstrated that Cyt c did not degrade upon treatment with denaturing agents in the presence or absence of ATP and [Cho][Sal] (Figure 7d), thus, confirming that the variation in the activity was due to the changes in the structural conformation of the protein.

### Nano-structured hydrotrope-caged Cyt c is resistant to protease digestion

Trypsin, being a proteolytic enzyme, hydrolyses the pep-tide bond at basic amino acids such as Arg and Lys residues (Figure 8a), thereby denaturing the target protein.^[40]^ Native and hydrotrope-stabilized Cyt c systems were incubated with 6 μM trypsin at 37°C for 24 hr. After the interaction, only 25% of activity was retained for native Cyt c, whereas the hydrotropic systems showed excellent performance and maintained Cyt c resistant to the ac-tion of trypsin (Figure 8b). In the presence of ATP, Cyt c lost only ∼2-fold activity compared to activity at the 0 hour (∼700% retained), whereas IL and ATP+IL systems retained ∼61- and ∼67-fold relative activity, respectively. This excellent performance against protease digestion can be deduced by close examination of the interactions of ATP and IL with Cyt c. H-bonding analysis revealed that ATP has a higher tendency to interact with Lys, Thr, and Gln at both 26.85°C and 90°C temperatures, thus suggesting the efficacy of ATP to impart stability to Cyt c (Figures 6d-6f and Figure S12). Additionally, IL also greatly affected the dynamic nature by interacting with a wide range of residues. The affinity of ATP and IL towards Lys may be helpful to protect Cyt c by hindering the degradation mechanism of trypsin. The electrostatic and H-bond interactions that occur when Cyt c is nanostructured with ATP completely safeguard the protein from trypsin degradation (Figure 8a). This was further shown by non-reducing SDS-PAGE analysis of the Cyt c before and after digestion with trypsin (Figure 8c). Most of the native Cyt c in the presence of tryp-sin was found degraded (lane 2, Figure 8c). However, the protein was intact when it was digested in the presence of ATP, IL and a combination of both. From UV-vis spectra (Figure S16), the Q band is lost in almost all the cases (except with ATP) after incubation with trypsin. However, only marginal changes were observed in the Soret band of Cyt c+ATP, and Cyt c+ATP+IL systems, indicating the stability induced by the nano-structured hydrotropes.

**Figure 8:**
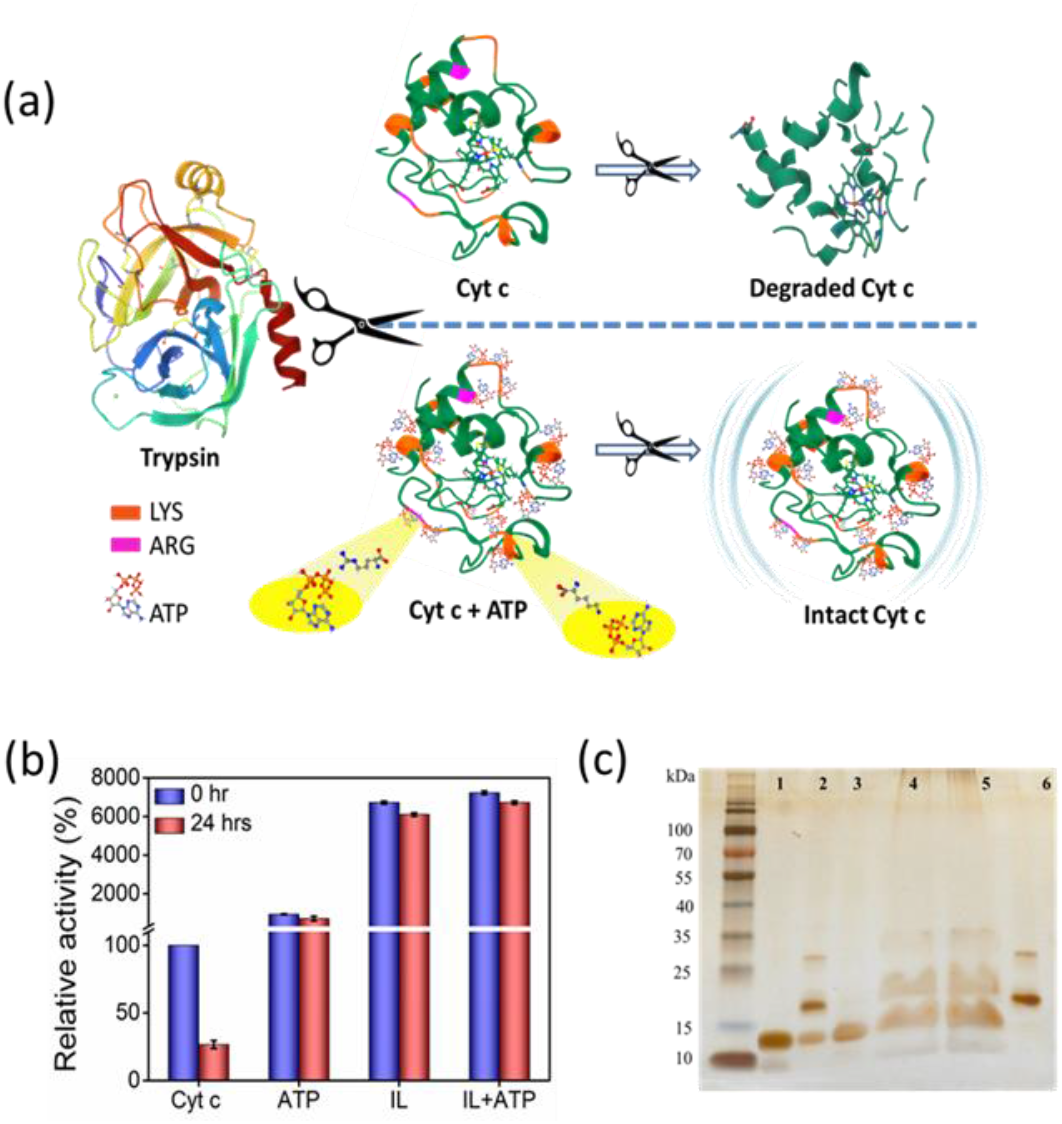
(a) Plausible mechanism of Cyt c digestion in the presence and absence of nano-structured hydrotropes. (b) Effect of pro-tease digestion (6 μM trypsin) on peroxidase activity of Cyt c. (c) SDS-PAGE of native Cyt c (lane 1), Cyt c incubated with trypsin (lane 2), Cyt c + ATP incubated with trypsin (lane 3), Cyt c + [Cho][Sal] incubated with trypsin (lane 4), Cyt C + ATP + [Cho][Sal] incubated with trypsin (lane 5), only trypsin (lane 6). In all cases, the concentration of ATP and IL was 5 mM and 300 mg/mL, respectively.

**Figure 9:**
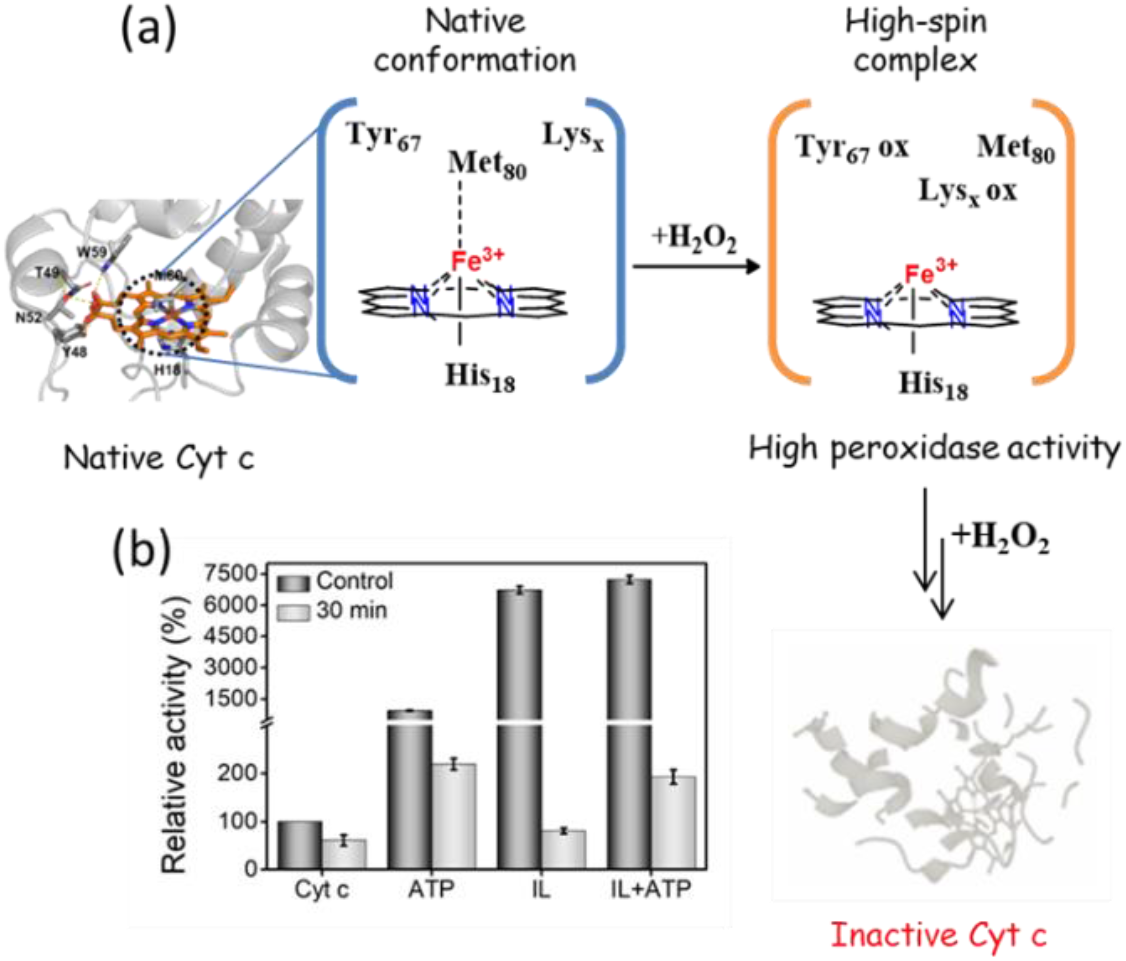
(a) Schematic mechanism showing H_2_O_2_ induced structural degradation of Cyt c. (b) Effect of H_2_O_2_ on the peroxidase activity of Cyt c in the presence of different hydrotropes.

### Nano-structured hydrotrope-caged Cyt c reaction to oxidative stress

Unlike other stress conditions where the peptide back-bone is attacked, H_2_O_2_ initially targets the catalytic site, followed by a breakdown of peptide bonds. Thus, even though H_2_O_2_ is essential for the peroxidase activity of Cyt c, upon prolonged exposure it causes severe impairment to the protein structure. In our earlier activity studies, 1 mM of H_2_O_2_ was added at the end, whereas for this study, Cyt c was incubated with H_2_O_2_ for 30 mins, and the substrate was added afterwards. The activity of bare Cyt c following the induction of oxidative stress was found to be reduced to approximately 60% of its original activity. Conversely, in the presence of ATP, ∼219% of activity was retained. This observation can be correlated with the earlier findings in which ATP was found to prevent the oxidation of Cyt c from the Cyt c oxidase enzyme in the mitochondria by hampering the electron flow rate.^[41-42]^ Alternatively, there was a severe drop in the activity of Cyt c from 6700% to just 80% in the presence of IL. These results are not astonishing since the salicylate counterpart is known to enhance the formation of ROS.^[22]^ However, the presence of ATP was found to prevent the oxidation and Cyt c from IL-induced oxidative stress and demonstrated 192% activity. These findings demonstrate the stabilizing effect of nano-structured ATP surrounding Cyt c.

## Conclusion

In summary, this study systematically demonstrated the robustness of ATP and IL-based mixed nano-structured hydrotropes and their utility in the improvement of protein packaging in extreme conditions. From the Mb solubility data, it was shown that the hydrotropic nature of ATP accelerated with the addition of a co-hydrotrope like [Cho][Sal] and vice-versa. Through RDF, DLS, ζ-potential, and UV-vis data, we provided convincing data that ATP forms oligomeric nano-structures at concentrations above 2.5 mM around Cyt c, which can be augmented further with the addition of [Cho][Sal]. A 9-fold increase in peroxidase activity with ATP, 67-fold with IL, and 72-fold with IL+ATP suggested a ‘partial reversible unfolding-electrostatic stabilization’ mechanism. Reversible binding of ATP and IL with the Ω 40-54 residue loop region of Cyt c showed higher stability at 26.85°C. Moreover, specific binding of nano-structured hydrotropes with Cyt c to Lys and Arg residues through H-bonding and polar interactions with Ω 70-85 region, presented an exceptionally high thermal tolerance with 80-fold activity even at 90oC. Because of such binding specificity, the structure and activity of nano-confined Cyt c were found to be retained against protease digestion, which explicitly cleaves at Lys and Arg residues. Furthermore, ATP and IL-based nano-structured hydrotropes showed efficacy in retaining the functional integrity of Cyt c even in the presence of chemical denaturants like urea and GuHCl, which suggested the suitability of these solvent manipulation strategies for industrial bio-catalysis. Through oxidative stress studies, a stark observation was made where [Cho][Sal] was found to enhance ROS formation, which could be efficiently subdued by adding ATP. These results have the potential for counteracting oxidative stress inside the living cells. Thus, the novel strategy of protein confinement in nano-structured hydrotropes can find significant usage in protein packaging under biotic and abiotic stresses, high-temperature bio-catalysis, and cell protection against ROS.

## Supporting Information

Supporting Information - Details of materials and methods, experimental processes, and figures related to DLS, zeta potential, UV-vis, and computational results have been provided. The parameters and input files for MD simulations variants, the key restart files, representative conformations of metastable states, and raw outputs from analyses are freely available at https://doi.org/10.5281/zenodo.7602239.

## Acknowledgements

D.M. acknowledges SERB, India (EEQ/2021/000059), for the research grant. D.K.S. was a recipient of a scholarship pro-vided by POWER project POWR.03.02.00-00-I006/17. The computations were performed at the Poznan Supercomputing and Networking Center. D.M. and G.F. acknowledge the NANOPLANT project, which received funding from the European Union’s Horizon 2020 research and innovation program under grant agreement no. 856961.

