## Supporting information for "Nano-structured Hydrotrope-Caged Cytochrome c with Boosted Stability in Harsh Environments: A Molecular Insight"

---

[a] PB, Dr. SMS, Prof. SKN, Dr. DM
Centre for Nano and Material Sciences
Jain (Deemed-to-be University)
Jain Global Campus, Kanakapura, Bangalore, Karnataka, India – 562112


[b] DKS, Prof. JB
Laboratory of Biomolecular Interactions and Transport, Department of Gene Expression, Institute of Molecular Biology and Biotechnology, Faculty of Biology
Adam Mickiewicz University
Uniwersytetu Poznanskiego 6, 61-614 Poznan, Poland


[c] DKS, Prof. JB
International Institute of Molecular and Cell Biology in Warsaw
Ks Trojdena 4, 02-109 Warsaw, Poland

[d] Dr. MB, Dr. VL, Prof. GF, Dr. DM
Institute of Plant Genetics (IPG)
Polish Academy of Sciences
Strzeszyńska 34, 60-479 Poznań, Poland


*Corresponding authors

[+] These authors have contributed equally to this work.

| Content | Page |
| --- | --- |
| Experimental section | 3-5 |
| Figure S1. Depiction of non-covalent interactions between hydrotropic molecules | 6 |
| Figure S2. Concentration dependant Zeta- potential analysis of ATP | 7 |
| Figure S3. Zeta- potential analysis of [Chol][Sal] IL | 8 |
| Figure S4. Dynamic light scattering data of ATP in presence of IL | 8 |
| Figure S5. Distribution Functions of nanostructured solvents | 9 |
| Figure S6. Zeta potential of ATP with 2μM of Cyt c | 10 |
| Figure S7. DLS showing nanostructuring of ATP and IL around Cyt c | 10 |
| Figure S8. UV-Vis spectra of Cyt c in presence of optimized concentrations of hydrotropes | 11 |
| Figure S9. Characterization of open and closed metastable states at 26.85 °C | 12 |
| Figure S10. Top ranked H-bonding interactive residues of Cyt c with hydrotropes at 26.85 °C | 12 |
| Figure S11. H-bonding percentages of protein residues at 26.85 °C | 13 |
| Figure S12. RMSD, RMSF and Markov Models of Cyt c in presence of hydrotropes at 90 °C | 14 |
| Figure S13. Characterization of metastable states of Cyt c from adaptive simulation data at 90 °C | 15 |
| Figure S14. Adaptive simulation data showing percentage of H-bonding at 90 °C | 16 |
| Figure S15. UV-Vis spectra of Cyt c and nanostructured systems in presence of Urea | 17 |
| Figure S16. UV-Vis spectra of Cyt c and nanostructured systems in presence of Trypsin | 17 |
| Figure S17. Spectral separation data of metastable states at 26.85 °C | 18 |
| Figure S18. Spectral separation data of metastable states at 90 °C | 19 |
| Figure S19. Data representing timescale separations of slow processes at 26.85 °C | 20 |
| Figure S20. Data representing timescale separations of slow processes at 90 °C | 21 |
| References | 22 |

### EXPERIMENTAL SECTION

***Materials**:* Cytochrome c (Equine heart) with >95% purity, Myoglobin (Equine heart) with >90% purity, 80% w/w aqueous solution of choline bicarbonate and 2,2′-azino-bis(3-ethylbenzothiazoline-6 sulfonic acid) diammonium salt (ABTS) with >98% purity were purchased from Sigma-Aldrich, USA. Disodium salt of adenosine 5'-phosphate (ATP), salicylic acid with purity 99%, urea (with 99% purity) and trypsin 1:250 were procured from S D Fine-Chem Ltd., Mumbai, India. 30% (w/w) in $H_2O$ Hydrogen peroxide ($H_2O_2$) and guanidium hydrochloride (GuHCl) were purchased from Avra synthesis Pvt. Ltd., Hyderabad, India. All the chemicals are used without any further purification and laboratory produced double distilled water was used throughout the process.

***Synthesis of choline salicylate ([Cho][Sal]) IL***: Synthesis of choline salicylate is carried out by following an established procedure.[1] Weighed 1:1 mole ratio of choline bicarbonate and salicylic acid are transferred to 100 mL round bottom flask with methanol as the solvent. Reaction mixture is kept for continuous stirring at 70°C for 12 hours. After the completion of reaction, solvent remaining is removed using rota evaporator. The IL obtained is then washed with ethyl acetate followed by bubbling with $N_2$ gas with simultaneous removal of trace water by vacuum.

***Solubility study of myoglobin***: Solubility studies of Equine heart Myoglobin in aqueous and hydrotropic media (ATP, IL and IL+ATP) were carried out at the pH 7 by increasing the amount of Mb and keeping the concentrations of ATP and IL constant. The transmittance of the respective solutions were recorded at various time intervals depending on the complete solubility of Mb. %Transmittance value at 530 nm was taken into consideration, since absorbance or transmittance% at this wavelength depends only on concentration of Mb and not affected by presence of any other molecular species used in this experiment.

***Characterization techniques**:* UV-Vis absorbance spectra and activity of the Cyt c in the absence and presence of ATP and IL+ATP were recorded using a Shimadzu UV-1900 spectrophotometer with the highest resolution of 1 nm using matched 1 cm path length quartz cuvettes. For absorbance spectra, the scans were done in the whole UV-Vis range of 250-800 nm. Each sample spectrum was collected by taking Cyt c (10 µM) and hydrotropic systems. The average of three spectra after eliminating the relevant blank from the tentative spectrum was considered for further analysis. DLS and Zeta potential studies were carried out using Anton Paar Litesizer 500 device in which prior to the analysis, the samplings were made similar to that of UV-Vis spectra. In addition, the samples were filtered through Moxcare Nexflo syringe filter of 0.22 µm pore size. DLS analysis was done in the quartz cuvette of 1cm path length and Omega cuvette Mat. No. 225288 was used for zeta potential. All the zeta potential studies were recorded in the protein mode with 200 cycles of run.

***SDS-PAGE analysis***: The role of ATP and [Cho][Sal] against different stresses such as high temperature, urea and trypsin digestion on structural integrity of Cyt c were analyzed by non-reducing SDS PAGE. 2 µM Cyt c was incubated with 5 mM ATP and 1.2 M [Cho][Sal] IL separately and in combination of both and were subjected to various stresses viz. high temperature (100°C for 10 min), 8M urea denaturation (30 min at room temperature), and trypsin digestion (24 h at 37°C). The samples were then analyzed by loading 250 ng equivalent Cyt c was run on non-reducing 12% Mini-Protean TGX Precast gels (BioRad), silver stained and imaged.

***Peroxidase activity of Cyt c at room temperature and under harsh conditions***: For activity profiling, a standard ABTS assay was employed where ABTS (3 mM) is oxidized to $ABTS^{+\cdot}$ in the presence of $H_2O_2$ (1 mM) and catalyzed by Cyt c (2 µM). All chemicals were kept at 25°C for an hour, followed by the incubation of Cyt c with different concentrations of ATP, IL, and IL+ATP for 10 mins. With the addition of ABTS and peroxide, the green color of ABTS intensifies due to the formation of cationic radicals, and variation of absorbance at 420 nm was monitored for 60 seconds with 1-sec intervals, resulting in a straight line. For the temperature studies, Cyt c alone, or incubated with hydrotropic systems were allowed to attain room temperature after subsequent heating (from 25-110°C for 30 mins). For chemical denaturants, 8 M of urea and 6 M of GuHCl were added to the native and pre-incubated catalytic systems, and the activity was checked at 30 and 15 mins, respectively. To profile the activity of Cyt c in the presence of a biological denaturant, the protein with and without hydrotropes was further incubated with 6 µM of trypsin at 37°C for 24 hrs. Finally, for oxidative stress examination, all systems were incubated with $H_2O_2$ for 15 mins, followed by the addition of ABTS. For all the activity studies, the slope of the curve obtained for native Cyt c was taken as 100%, and the activity of other systems was calculated relative to this.

***System setup and Molecular Dynamics simulation***: The input model was based on the crystallographic structure of bovine heart Cyt c (PDB code: 2B4Z) at 1.5 Å resolution. The force field parameters for the heme cofactor were produced following a literature search for which the geometry optimization was done using the Gaussian 16 program.[2] All the simulations were established using the AMBER 18 package,[3] and the parameters for ATP were retrieved from the AMBER parameter database.[4] Structures of [Cho][Sal] were obtained from the PubChem database (CID 54686350), and the force field parameters were generated using RESP-A1 charge model using a multi-orientational RESP fitting procedure integrated in the RESP ESP web-server and General Amber Force Field (GAFF2) parameters. [5,6] Four molecular systems were prepared, keeping the protein Cyt c in water, ATP, IL, and both ATP and IL. All molecular systems were solvated in a cubic box with a side length of 10 nm prepared using a PACKMOL package. [7] The number of ATP molecules corresponds to the concentration of 5 mM, which is 3 ATP molecules, and the number of IL corresponds to 300 mg/mL with a total of 350 molecules of [Cho][Sal]. The protein was modeled using the AMBER ff19SB force field.[8] The protein structure was further protonated using H++ web server pH 6.8.[9] The protein complexes were solvated using OPC water models and then neutralized with counter ions ($Na^+$ and $Cl^-$) to achieve the ionic strength of 0.1 M. The systems were then minimized using PMEMD and PMEMD.CUDA modules[10] of AMBER 18 and equilibrated under constant pressure and temperature (NPT) conditions at 1 atm and 300 K using the Langevin thermostat and periodic boundary conditions using the particle mesh Ewald method[11] with 4 fs time-steps enabled by SHAKE and hydrogen mass repartitioning algorithms.[12] Using the steepest descent minimization method, the systems were minimized for several rounds with decreasing harmonic restraints. The following steps included continued heating of the systems from 200 K to 300 K with restraints on only the heavy atoms for 1 ns, followed by NPT equilibration with backbone restraints for 1 ns. Finally, the production run was performed using several rounds of unrestrained NPT simulations with a frame saving rate of 200 ps and total sampling time of 1 ns at constant temperature and pressure using the weak-coupling barostat and thermostat.[13] The final configurations of protein in different solvent environments were adopted for simulations at room temperature (300 K; 26.85$^{o}$C) and higher temperatures (363.15 K; 90$^{o}$C). The last snapshots from the production run were used for the initial input structure for adaptive sampling simulations using the High-Throughput Molecular Dynamics (HTMD) package.[14] The dihedral angle of the protein backbone was defined as the guiding matrix in adaptive sampling simulations. A total of 10 epochs were sampled, with each epoch consisting of 10 individual simulations of 100 ns, totaling 10 μs. All the trajectories were analyzed using the cpptraj program and its Python front-end, pytraj.[15] Further classical MD of 200 ns lengths were performed for solvent systems of ATP, IL, and both ATP and IL without protein.

***Markov state models*:** The Markov state models (MSM) were obtained using the functionalities of the pyEMMA program implemented in the HTMD protocol.[16] The high dimensionality space was reduced using time-lagged independent component analysis (TICA). The models were generated, and the number of metastable states was selected based on the spectral separation analysis (Figures S17 & S18). A lag time of 40 ns was chosen to estimate the final Markov model (Figures S19 & S20).

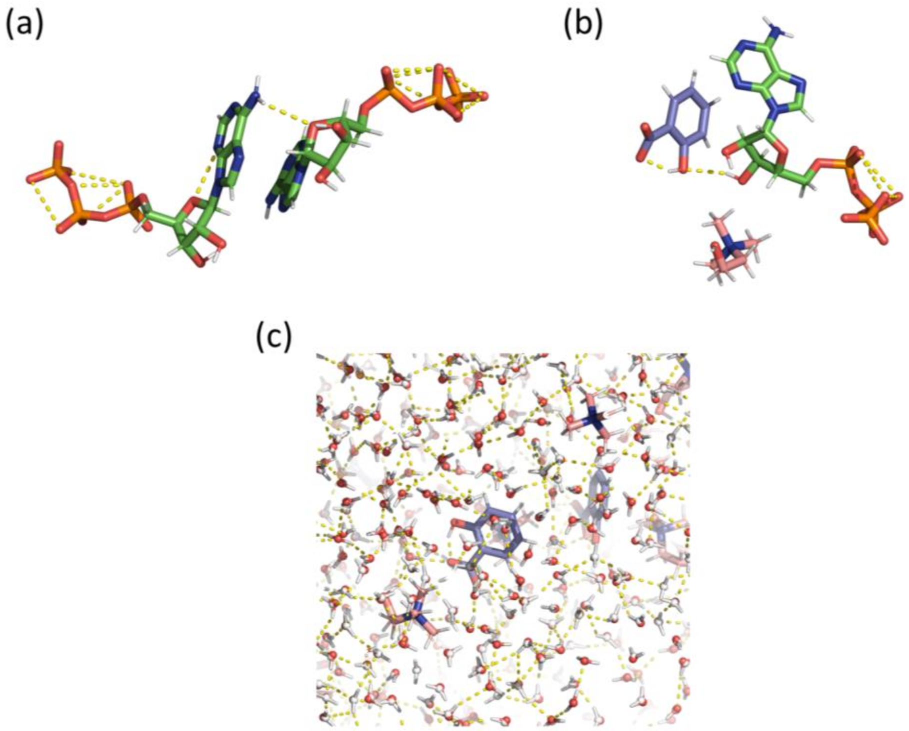


**Figure S1**. (a) Typical non-covalent interactions of two ATP molecules. (b) Representative non-covalent interactions of IL+ATP. (c) Non-covalent interactions with water molecules with IL (Cholinium in pink and salicylate in blue).

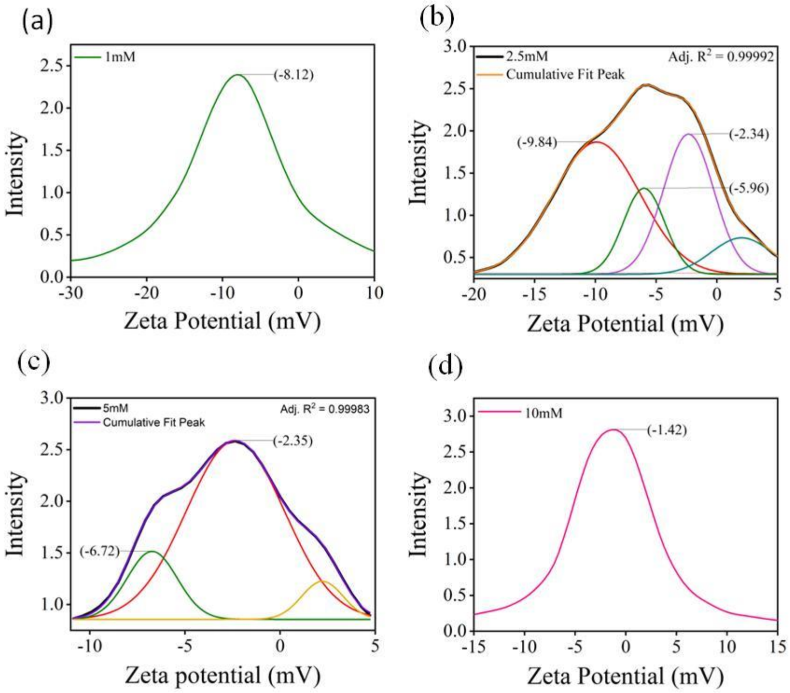


**Figure S2**: Concentration dependent zeta potential analysis of ATP with corresponding distribution peaks. (a) At 1 mM concentration, showing non-aggregated free form; (b) At 2.5 mM, deconvoluted peaks showing onset of nanostructuring and presence of free form; (c) at 5 mM concentration, showing the absence of free form and (d) single distribution peak at 10 mM, representing existence of only nanostructured state.

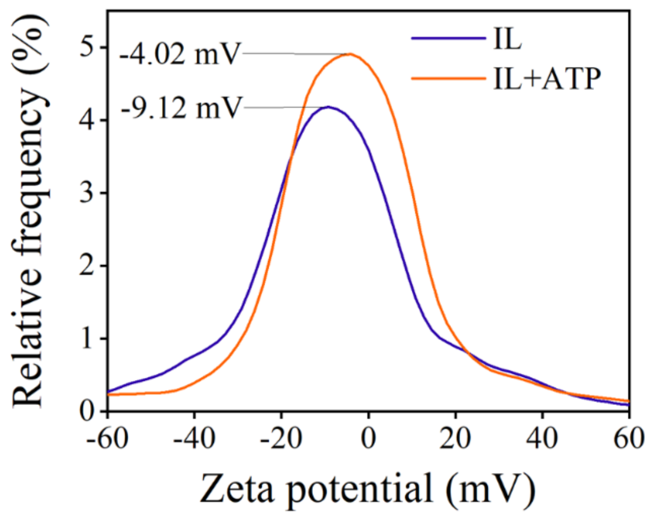


**Figure S3**: Zeta potential results of [Cho][Sal] IL and ATP+IL.

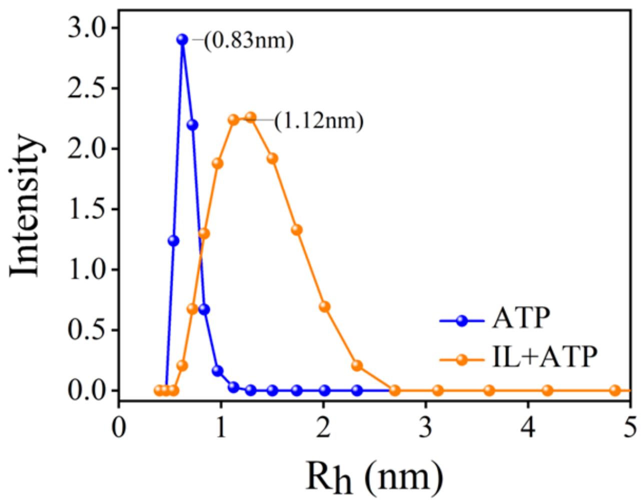


**Figure S4**: DLS analysis showing enhanced nano structuring of ATP in presence of [Cho][Sal] IL.

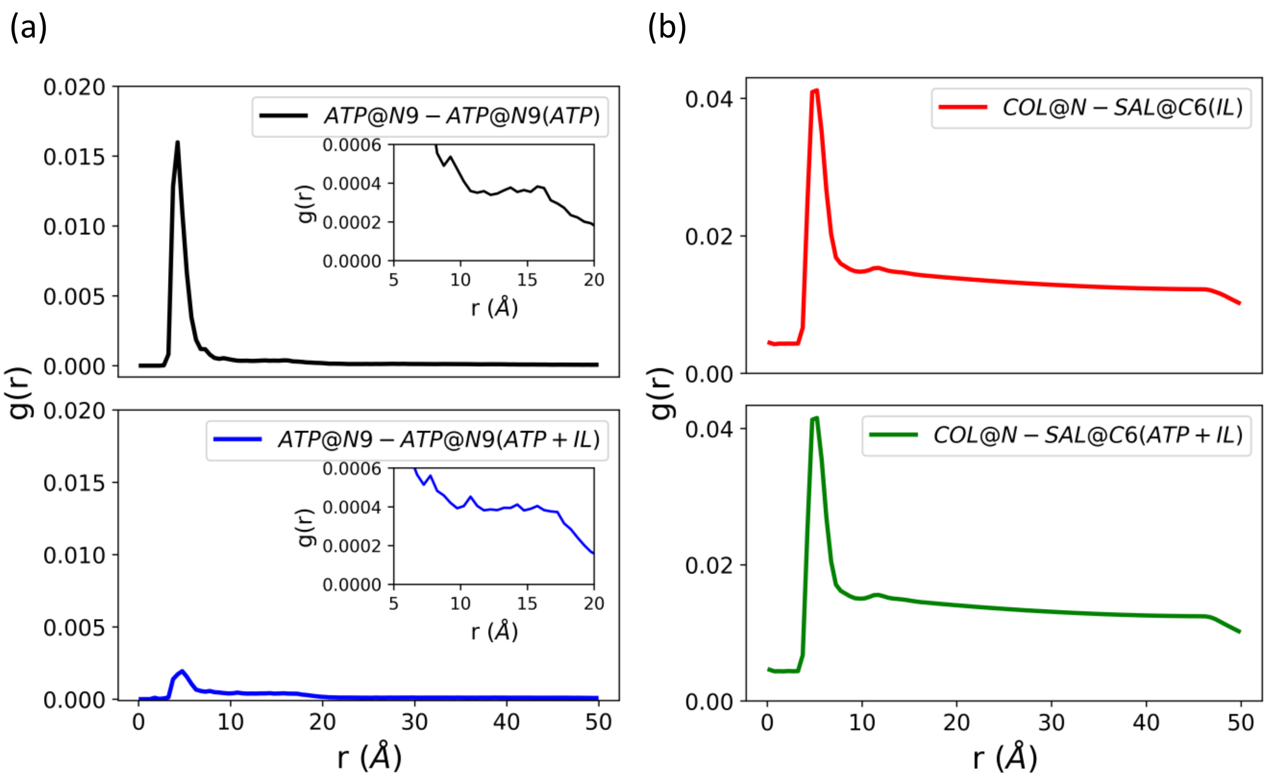


**Figure S5**. Radial Distribution Functions of nanostructured solvents from adaptive simulation of 10 µs of Cyt c in presence of (a) ATP (pair-wise distances of N9 atoms) in both ATP and ATP+IL molecular systems, (b) IL (pair-wise distances of N and C6 atoms) in both IL and ATP+IL molecular systems. All simulations were performed at 26.85°C.

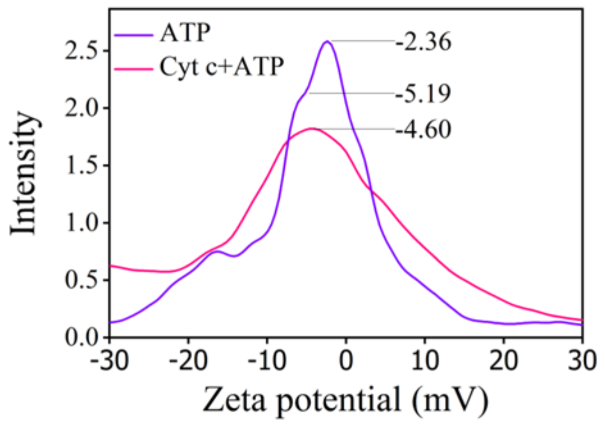


**Figure S6**: Zeta potential analysis of ATP with 2 μM of Cyt c.

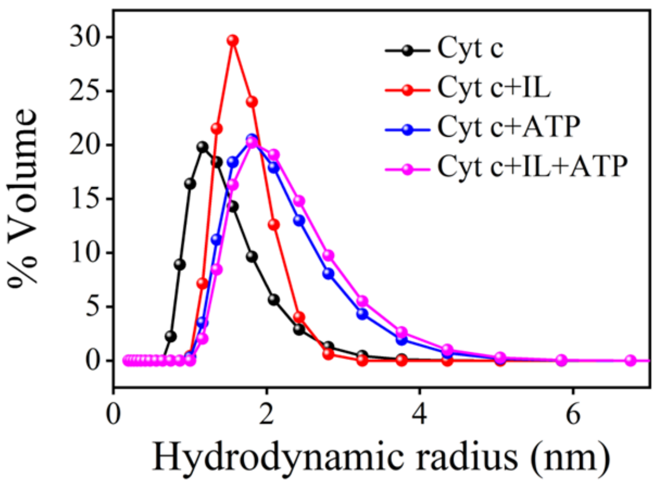


**Figure S7:** DLS distribution curve showing nanostructuring of ATP and IL around Cyt c.

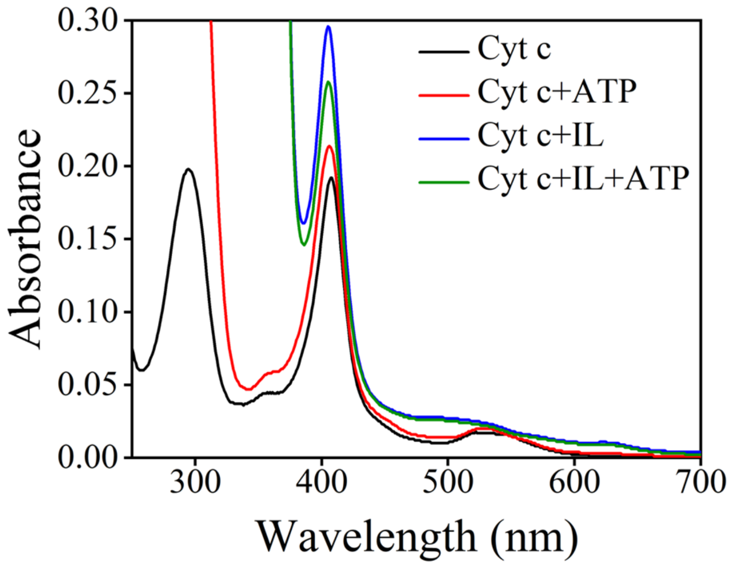


**Figure S8:** UV-Vis spectra of Cyt c in presence of optimized concentrations of ATP, IL and IL+ATP.

| | Avg. RMSD (Å) | RMSD (std.) | Avg. Rg (Å) | Rg (std.) | mfpton (ns) | std. | mfptoff (ns) | std. | Sample proportions |
|---|---|---|---|---|---|---|---|---|---|
| | | | | | **WAT** | | | | |
| model0 (open) | 5.55 | 0.70 | 14.00 | 0.37 | | | | | 0.30 |
| model1 * | 3.00 | 0.35 | 13.07 | 0.16 | 10.40 | 10.80 | 10.70 | 1.24 | 0.41 |
| model2 (close) | 2.30 | 0.27 | 12.87 | 0.12 | 53.30 | 9.79 | 917.00 | 1.24 | 0.28 |
| | | | | | **ATP** | | | | |
| model0 (open) | 5.09 | 0.54 | 13.66 | 0.24 | | | | | 0.10 |
| model1 * | 2.84 | 0.26 | 13.05 | 0.15 | 13.70 | 4.45 | 30.80 | 10.56 | 0.41 |
| model2 (close) | 2.12 | 0.20 | 12.81 | 0.10 | 109.00 | 9.88 | 746.00 | 10.46 | 0.49 |
| | | | | | **IL** | | | | |
| model0 (close) | 2.10 | 0.26 | 12.81 | 0.11 | 229.00 | 14.39 | 176.00 | 8.39 | 0.15 |
| model1 * | 3.25 | 0.49 | 13.18 | 0.19 | 130.00 | 16.90 | 24.80 | 8.45 | 0.34 |
| model2 (open) | 5.80 | 0.78 | 14.16 | 0.40 | | | | | 0.50 |
| | | | | | **ATP+ IL** | | | | |
| model0 (open) | 6.51 | 1.32 | 14.46 | 0.67 | | | | | 0.65 |
| model1 (close) | 2.49 | 0.37 | 12.94 | 0.16 | 82.00 | 5.11 | 236.00 | 2.47 | 0.35 |

*source = open; sink= close; * treated as source*

**Figure S9**: Characterization of open and closed metastable states from adaptive simulation data of 10µs for molecular systems of Cyt c in presence of ATP, IL, and ATP+IL. The mean fast passage time for association (open to close) and dissociation (close to open) are given as *mfpton* and *mfptoff* respectively. The average root-mean square deviation, radius of gyration is depicted *Avg. RMSD* and *Avg. Rg* respectively. The standard deviations are abbreviated as *std.* All simulations were performed at 26.85°C.

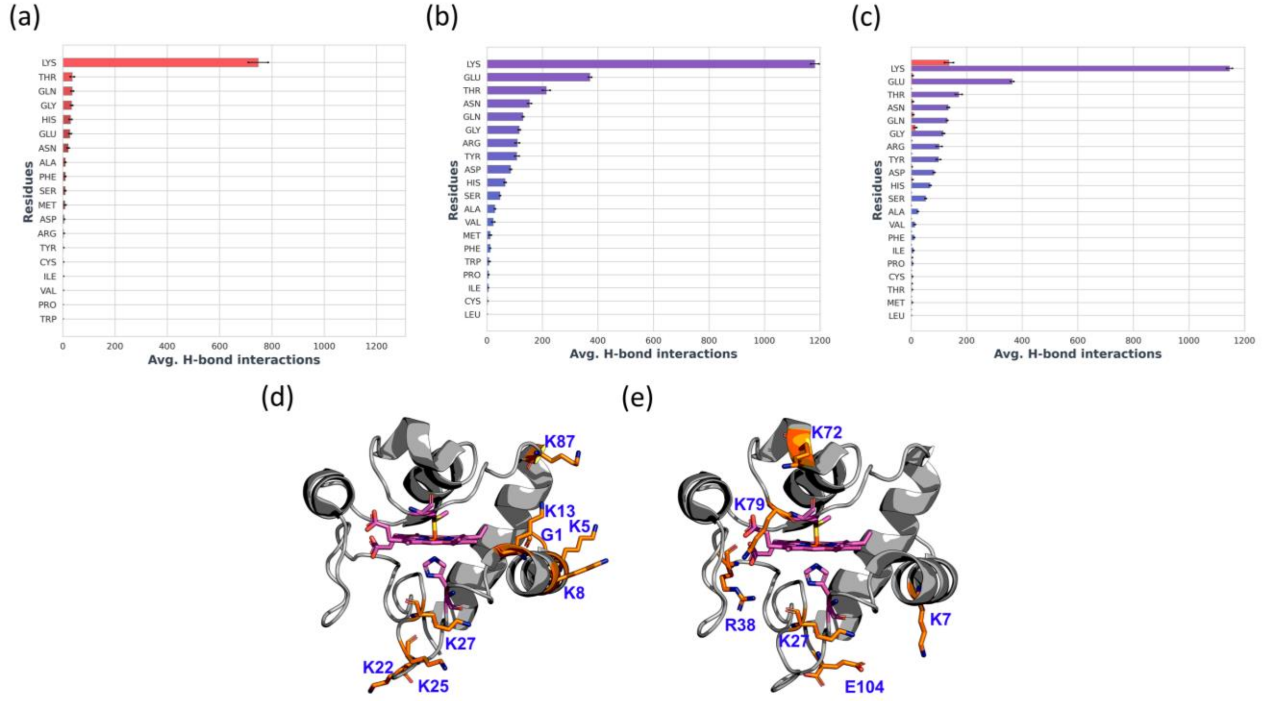


**Figure S10**: H-bonding interaction counts per protein residue types per simulation obtained from adaptive simulation data of 10µs for molecular systems of Cyt c in presence of (a) ATP, (b) IL, (c) ATP+IL. ATP interaction counts are shown red and IL in violet. The positions of top five interacting residues for ATP (from both ATP and ATP+IL systems) and IL (from both IL and ATP+IL) are shown in (d) and (e) respectively. All simulations were performed at 26.85°C.

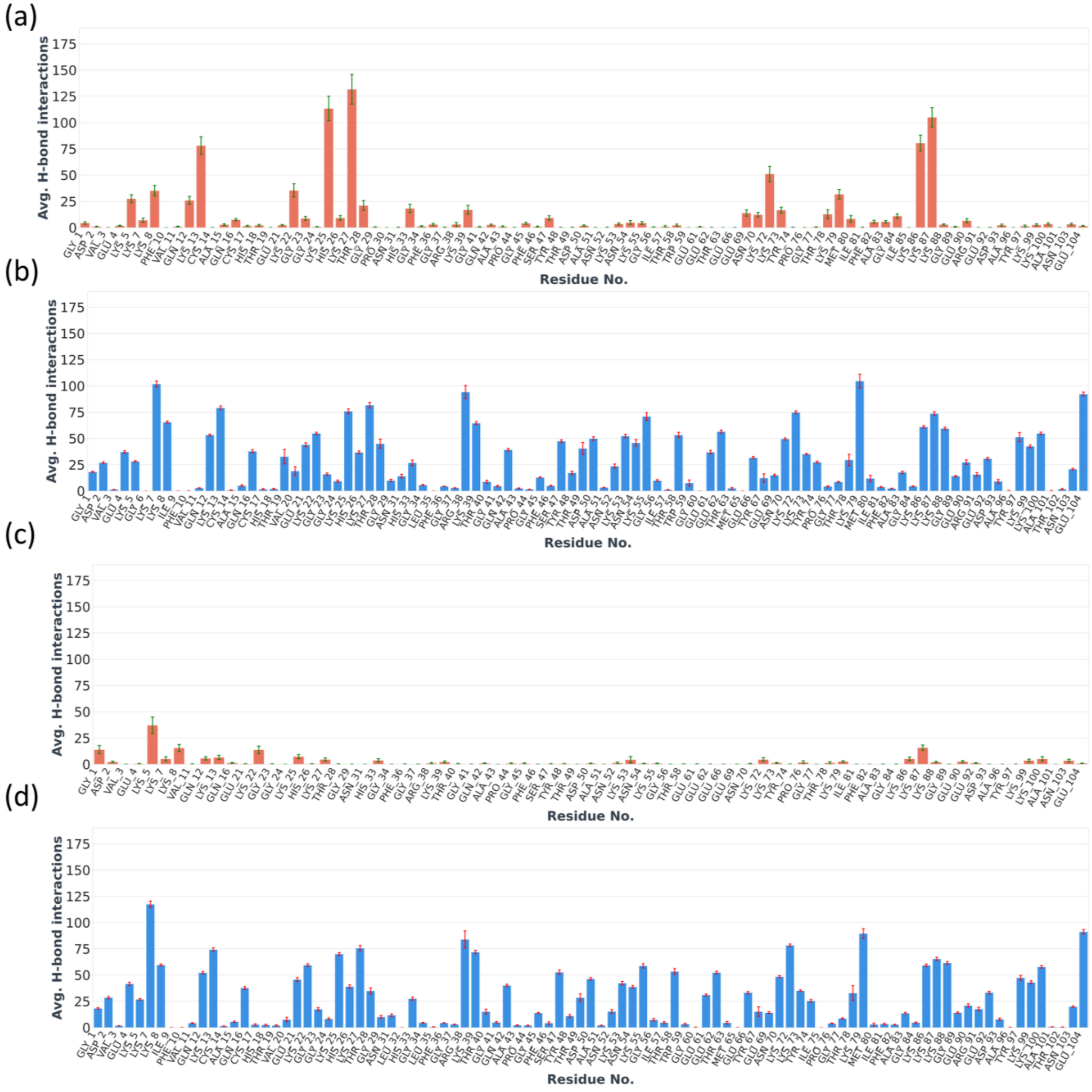


**Figure S11**: H-bonding percentages of protein residues obtained from adaptive simulation data of 10 µs for molecular systems of Cyt c in presence of (a) ATP, (b) IL, (c) ATP (ATP+IL) and (d) IL (ATP+IL). All simulations were performed at 26.85°C.

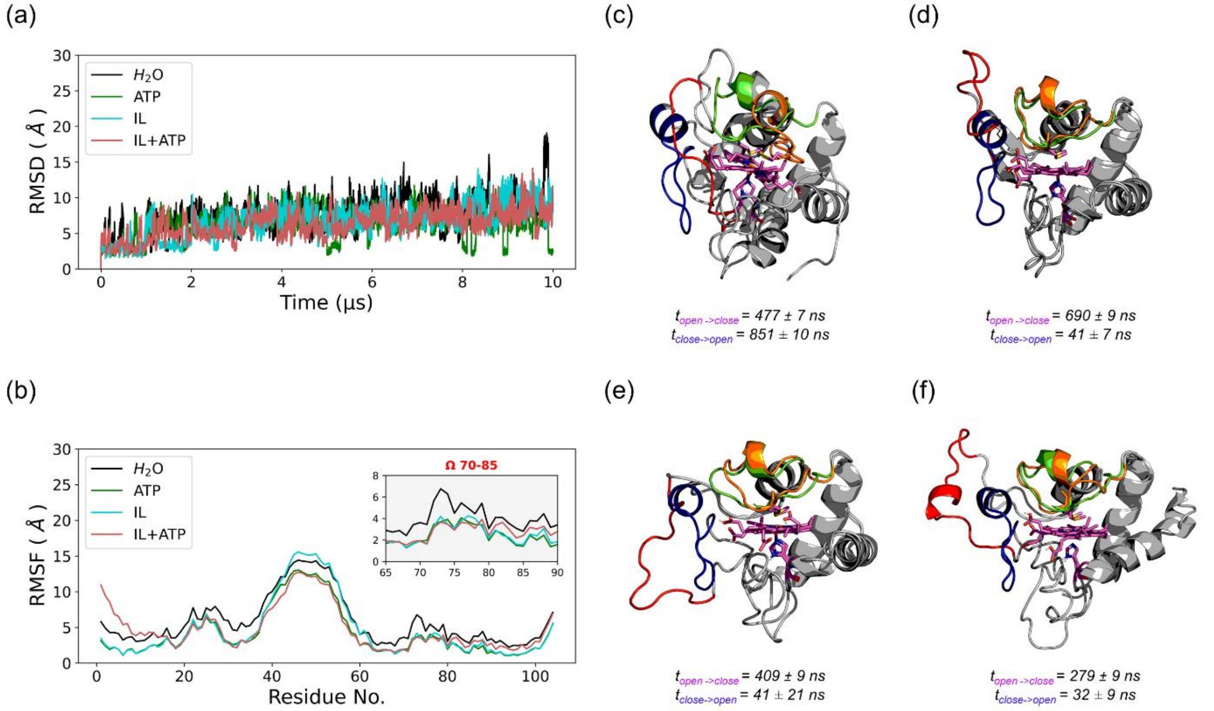


**Figure S12**: RMSD and RMSF of Cyt c shown in plot (a) and (b) calculated from 10μs of adaptive simulation. Markov models depicting open and closed metastable states with highly flexible region 40-54 highlighted for open(red) and close(blue) the metastable states as well as functional (green) and unstable (orange) states of region 70-85. The mean first passage time is shown for open to close and vise-versa for molecular systems (c) $H_2O$, (d) ATP, (e) IL and (f) both ATP+IL. All simulations were performed at 90°C.

| | *Avg. RMSD (Å)* | *RMSD (std.)* | *Avg. Rg (Å)* | *Rg (std.)* | *mfpton (ns)* | *std.* | *mfptoff (ns)* | *std.* | *Sample proportions* |
|---|---|---|---|---|---|---|---|---|---|
| | | | | | ***WAT*** | | | | |
| *model0 (open)* | *15.75* | *1.15* | *15.64* | *1.06* | | | | | *0.23* |
| *model1 (close)* | *2.91* | *0.36* | *13.01* | *0.15* | *477.00* | *6.69* | *851.00* | *9.42* | *0.09* |
| *model2 ** | *5.49* | *0.86* | *13.63* | *0.44* | *8.78* | *9.40* | *394.00* | *6.32* | *0.35* |
| *model3 ** | *8.69* | *1.26* | *14.22* | *0.65* | *35.40* | *9.46* | *448.00* | *2.23* | *0.33* |
| | | | | | ***ATP*** | | | | |
| *model0 (close)* | *2.58* | *0.42* | *12.93* | *0.14* | *604.00* | *9.16* | *41.10* | *7.26* | *0.08* |
| *model1 (open)* | *6.61* | *1.26* | *13.77* | *0.51* | | | | | *0.92* |
| | | | | | ***IL*** | | | | |
| *model0 (close)* | *3.20* | *0.52* | *13.15* | *0.22* | *409.00* | *9.01* | *41.70* | *20.99* | *0.10* |
| *model1 (open)* | *7.55* | *1.56* | *14.32* | *0.70* | | | | | *0.90* |
| | | | | | ***ATP+ IL*** | | | | |
| *model0 (close)* | *2.74* | *0.43* | *12.99* | *0.16* | *279.00* | *8.90* | *31.80* | *8.55* | *0.13* |
| *model1 ** | *5.15* | *0.85* | *13.66* | *0.38* | *227.00* | *6.62* | *5.75* | *8.67* | *0.33* |
| *model2 (open)* | *7.77* | *1.12* | *14.59* | *0.84* | | | | | *0.54* |
| | | | | | *source = open; sink= close; * treated as source* | | | | |

**Figure S13**: Characterization of open and closed metastable states from adaptive simulation data of 10μs for molecular systems of Cyt c in presence of ATP, IL, and ATP+IL. The mean fast passage time for association (open to close) and dissociation (close to open) are given as *mfpton* and *mfptoff* respectively. The average root-mean square deviation, radius of gyration is depicted *Avg. RMSD* and *Avg. Rg* respectively. The standard deviations are abbreviated as *std.* All simulations were performed at 90°C.

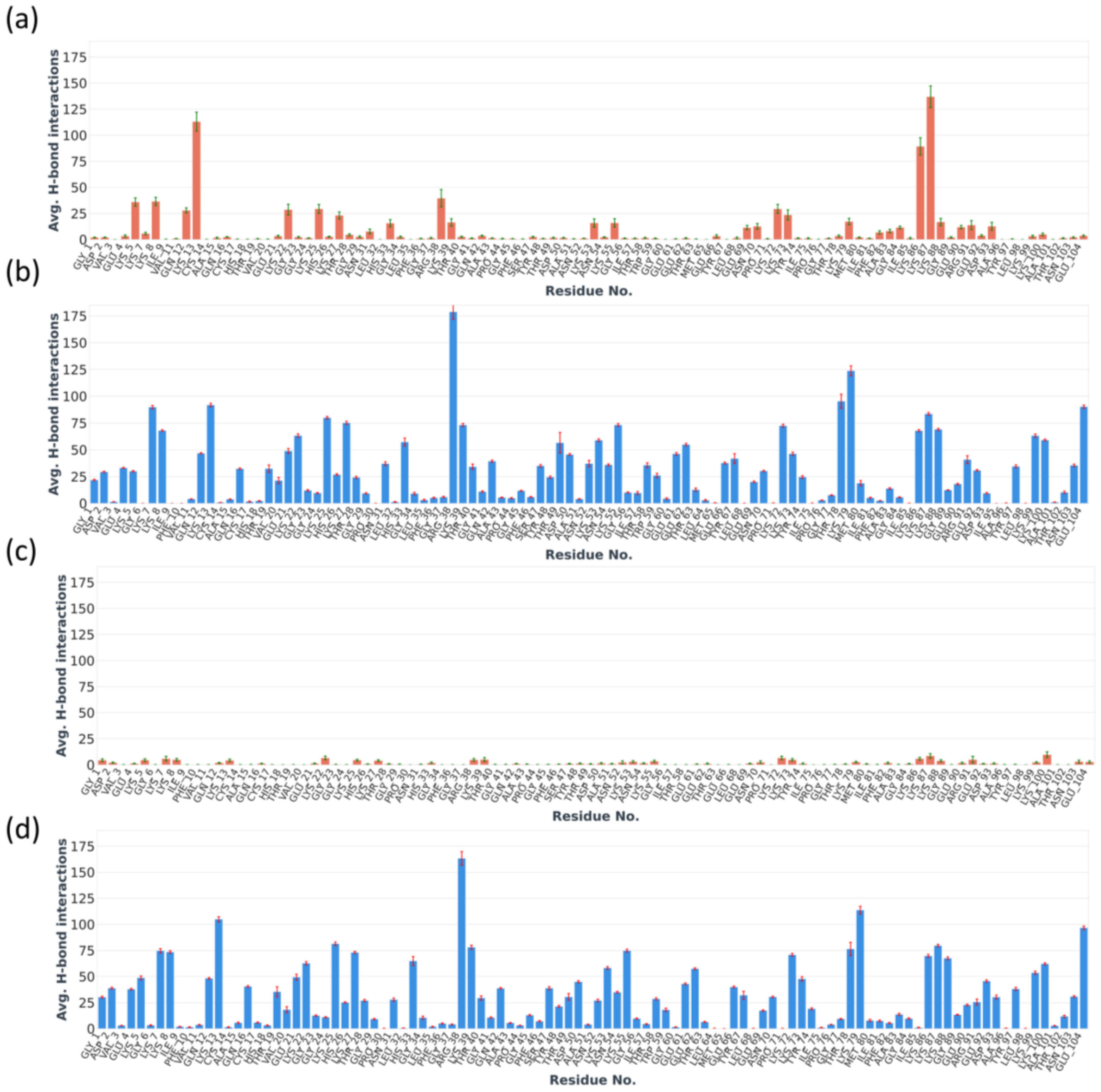


**Figure S14**: H-bonding percentages of protein residues obtained from adaptive simulation data of 10 μs for molecular systems of Cyt c in presence of (a) ATP, (b) IL, (c) ATP (ATP+IL) and (d) IL (ATP+IL). All simulations were performed at 90°C.

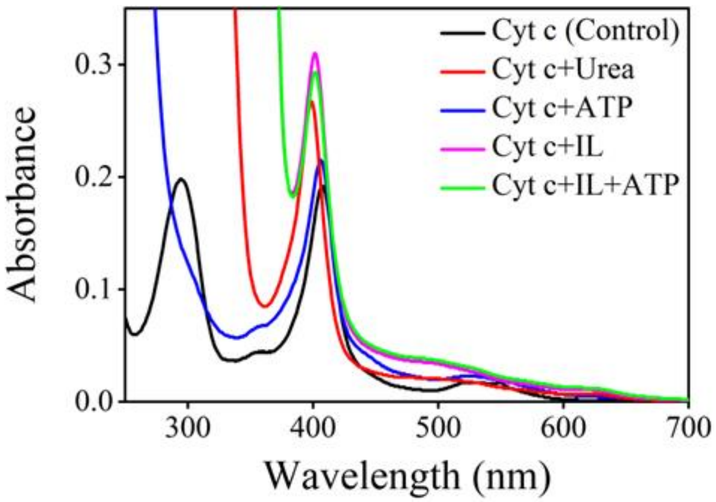


**Figure S15**: UV-vis spectra of Cyt c and nanostructured systems in presence of 8 M Urea.

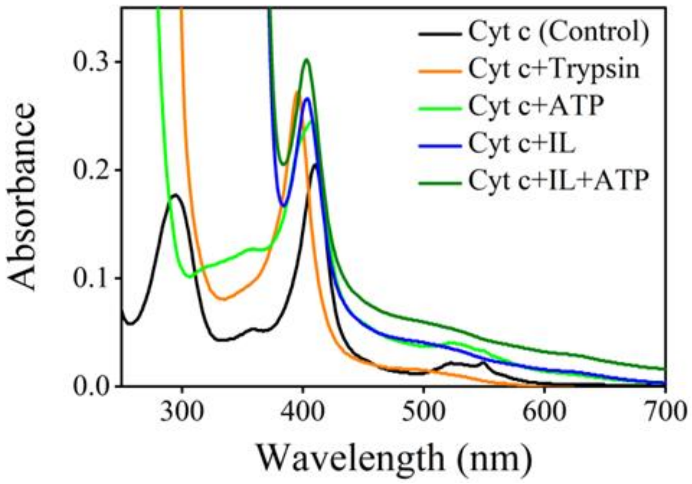


**Figure S16**: UV-Vis spectra of Cyt c and nanostructured systems in presence of Trypsin.

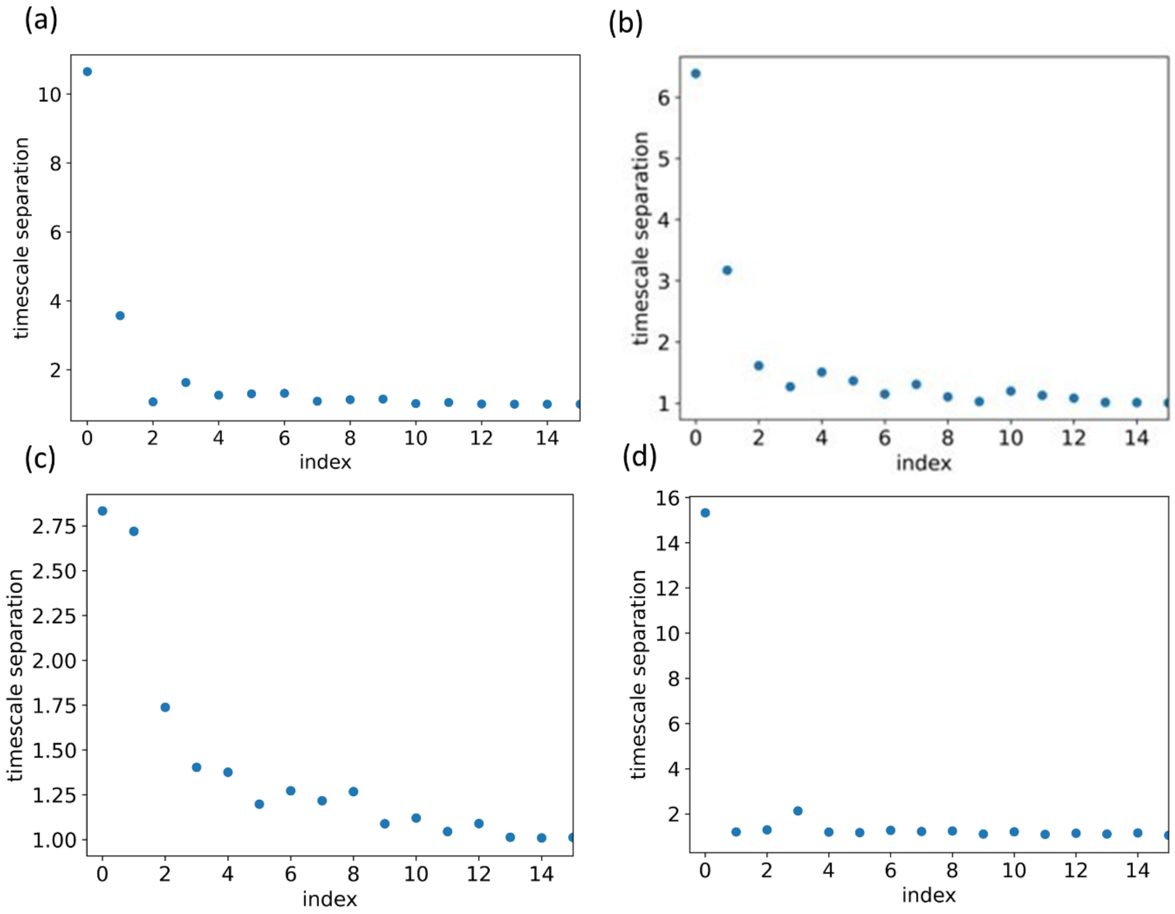


**Figure S17:** Spectral separations for selection of relevant metastable states obtained from adaptive simulation of Cyt c in presence of (a) $H_2O$, (b) ATP, (c) IL and (d) both ATP+IL. All simulations were performed at 26.85°C.

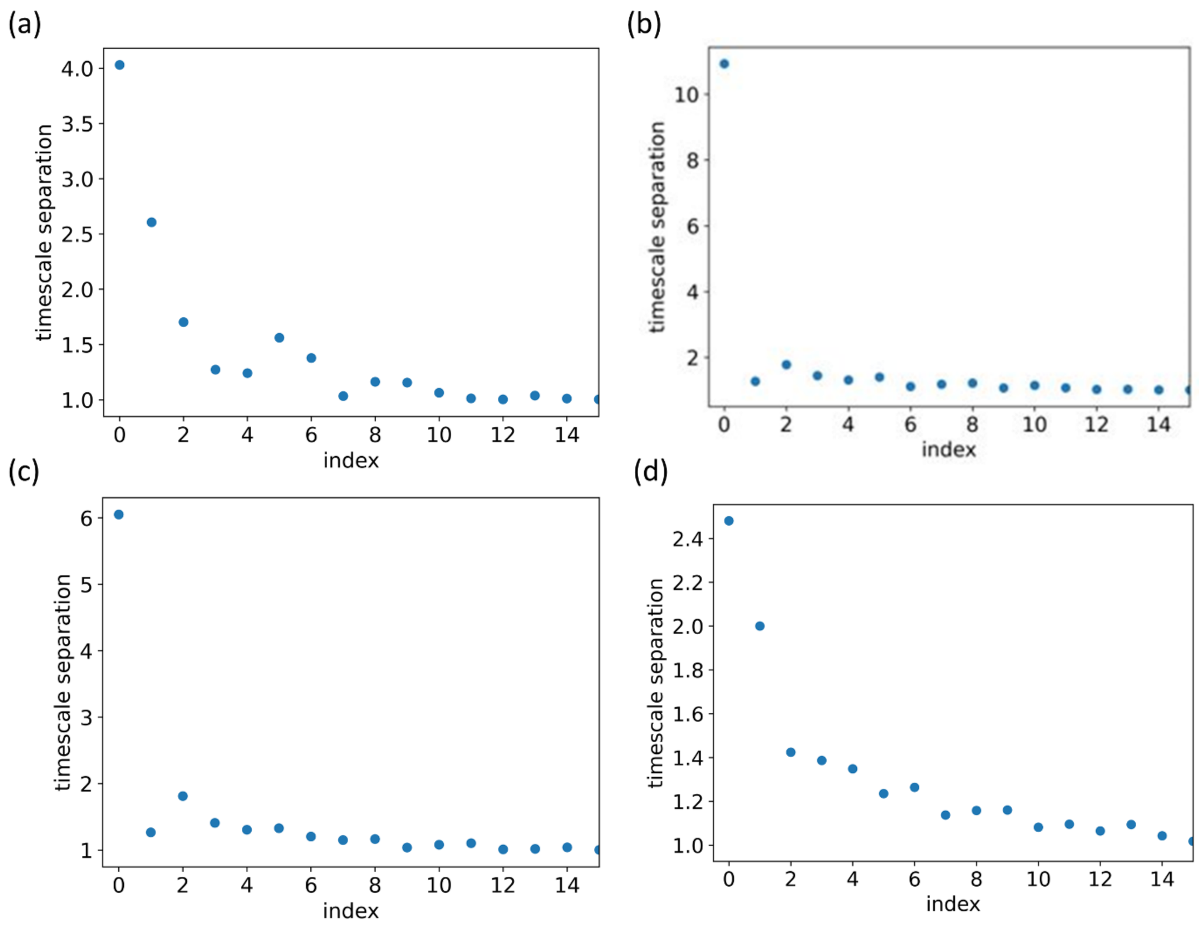


**Figure S18**: Spectral separations for selection of relevant metastable states obtained from adaptive simulation of Cyt c in presence of (a) $H_2O$, (b) ATP, (c) IL and (d) both ATP+IL. All simulations were performed at 90°C.

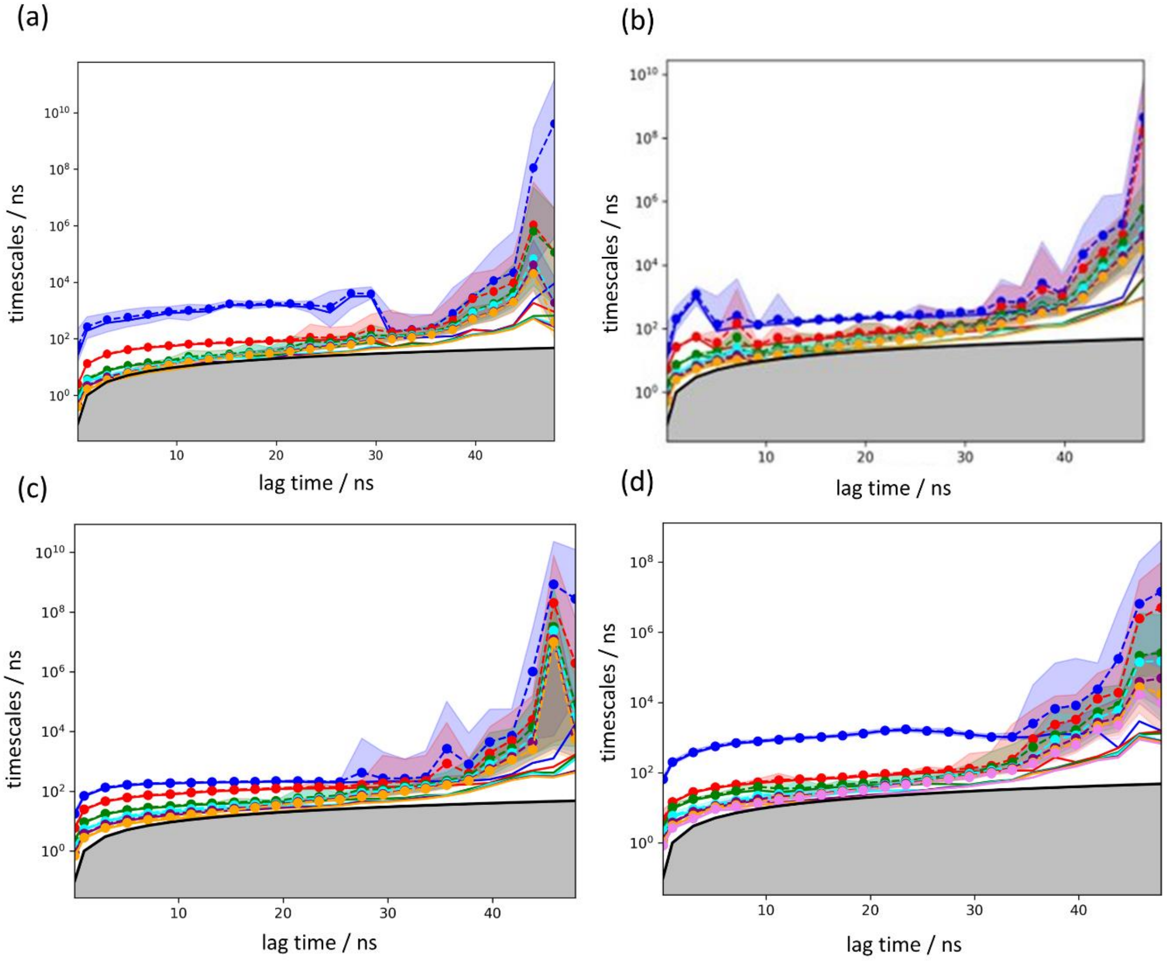


**Figure S19:** Implied time scales showing timescale separations of slow processes identified before building Markov models from adaptive simulation of Cyt c in presence of (a) $H_2O$, (b) ATP, (c) IL and (d) both ATP+IL. All simulations were performed at 26.85°C.

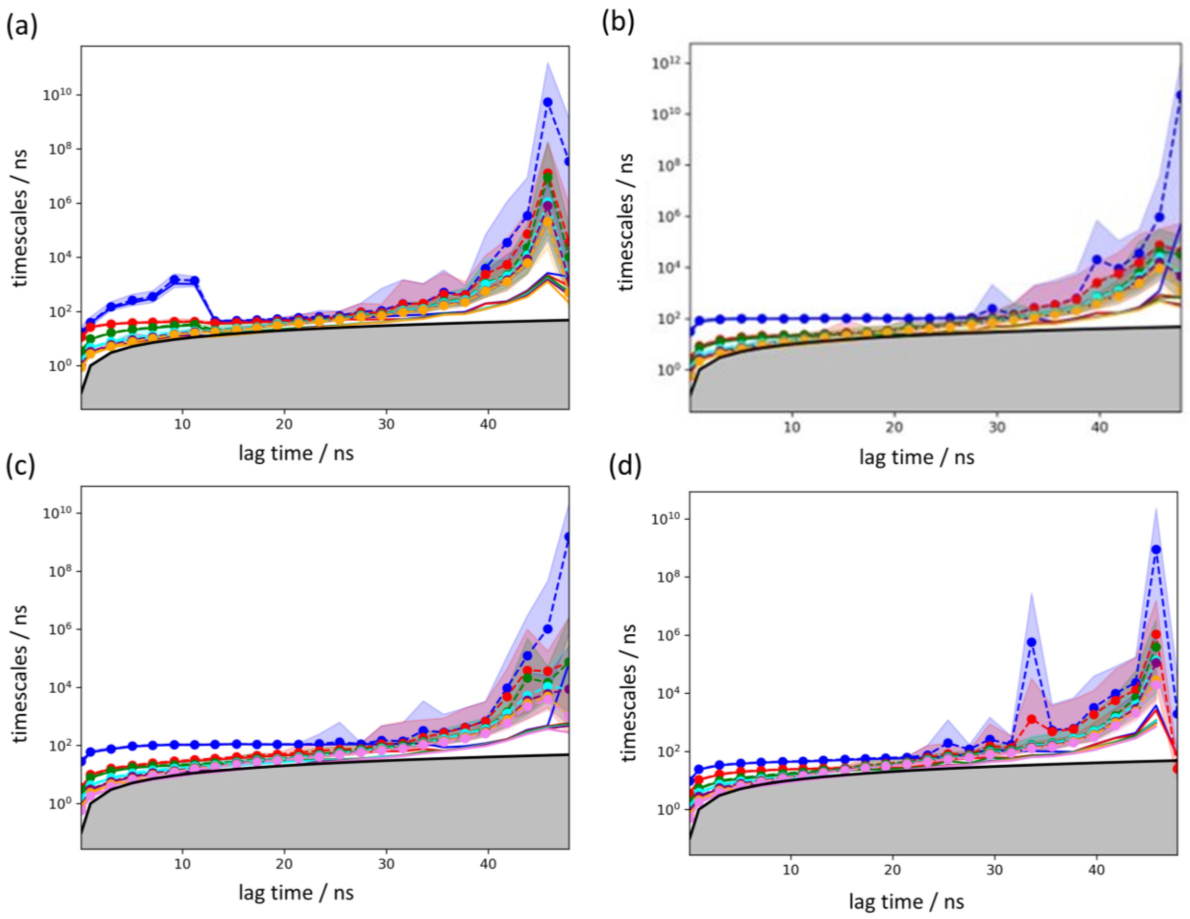


**Figure S20**: Implied time scales showing timescale separations of slow processes identified before building Markov models from adaptive simulation of Cyt c in presence of (a) $H_2O$, (b) ATP, (c) IL and (d) both ATP+IL. All simulations were performed at 90°C.
